# On the illusion of auxotrophy: *met15Δ* yeast cells can grow on inorganic sulfur thanks to the previously uncharacterized homocysteine synthase Yll058w

**DOI:** 10.1101/2022.01.19.476918

**Authors:** S. Branden Van Oss, Saurin Bipin Parikh, Nelson Castilho Coelho, Aaron Wacholder, Ivan Belashov, Sara Zdancewicz, Manuel Michaca, Jiazhen Xu, Yun Pyo Kang, Nathan P. Ward, Sang Jun Yoon, Katherine M. McCourt, Jake McKee, Trey Ideker, Andrew P. VanDemark, Gina M. DeNicola, Anne-Ruxandra Carvunis

## Abstract

Organisms must either synthesize or assimilate essential organic compounds to survive. The homocysteine synthase Met15 has been considered essential for inorganic sulfur assimilation in yeast since its discovery in the 1970s. As a result, *MET15* has served as a genetic marker for hundreds of experiments that play a foundational role in eukaryote genetics and systems biology. Nevertheless, we demonstrate here through structural and evolutionary modeling, *in vitro* kinetic assays, and genetic complementation, that an alternative homocysteine synthase encoded by the previously uncharacterized gene YLL058W enables cells lacking Met15 to assimilate enough inorganic sulfur for survival and proliferation. These cells however fail to grow in patches or liquid cultures unless provided with exogenous methionine or other organosulfurs. We show that this growth failure, which has historically justified the status of *MET15* as a classic auxotrophic marker, is largely explained by toxic accumulation of the gas hydrogen sulfide due to a metabolic bottleneck. When patched or cultured with a hydrogen sulfide chelator, and when propagated as colony grids, cells without Met15 assimilate inorganic sulfur and grow, and cells with Met15 achieve even higher yields. Thus, Met15 is not essential for inorganic sulfur assimilation in yeast. Instead, *MET15* is the first example of a yeast gene whose loss conditionally prevents growth in a manner that depends on local gas exchange. Our results have broad implications for investigations of sulfur metabolism, including studies of stress response, methionine restriction, and aging. More generally, our findings illustrate how unappreciated experimental variables can obfuscate biological discovery.

## INTRODUCTION

The metabolism of organic sulfur-containing compounds (“organosulfurs”) is critical for all domains of life. Plants and microbes, unlike animals, often synthesize organosulfurs from inorganic sulfates via the sulfate assimilation pathway (SAP) (1). Enzymes in the SAP reduce inorganic sulfates into sulfides and ultimately into organosulfurs. In yeast, the terminal step of the SAP generates homocysteine from O-acetyl-L-homoserine (OAH) and hydrogen sulfide (H_2_S). Homocysteine can then be converted either directly to methionine or indirectly to cysteine. These amino acids may be incorporated into nascent polypeptides; alternatively, they may be modified to produce other essential organosulfurs, including the “universal methyl donor” S-adenosylmethionine (SAM) and the important buffer against oxidative stress glutathione. Reactions downstream of homocysteine biosynthesis are reversible; thus all of these organosulfurs may be recycled back to homocysteine (1).

In *Saccharomyces cerevisiae*, Met15 (also referred to as Met17 and Met25) catalyzes homocysteine biosynthesis (2-4). The first precise and complete deletion of *MET15* (*met15*Δ*0*) in *S. cerevisiae* was made in 1998 (5). Many commonly used laboratory strains contain this deletion, including BY4741, which serves as the genetic background of the haploid *MAT***a** version of the yeast deletion collection that has been extensively utilized for functional genomics and other purposes (6, 7). The use of this deletion as an auxotrophic marker, and the corresponding assumption that *MET15* is required for growth in organosulfur-free media, has been applied to the interpretation of a wide range of studies (8-10). However, some evidence suggests that *MET15* may not be strictly required for growth without exogenous organosulfurs. For example, the study that first identified *MET15* (11) reported that the organosulfur requirement of *met15* mutants was alleviated at a critical methylmercury concentration. Papillae have also been seen when replica plating thick patches of *met15*Δ*0* cells to medium lacking organosulfurs (12).

Here, we demonstrate that *met15*Δ cells can synthesize organosulfurs from inorganic sulfates despite a long-held assumption to the contrary. We characterize a novel homocysteine synthase, Yll058w, and show that its activity enables *met15*Δ cells to grow in the absence of exogenous organosulfurs in certain experimental conditions. Specifically, stable growth is observed with automated colony transfer, but very low or no growth is observed when using traditional techniques such as thick patches or liquid cultures. We show that this appearance of auxotrophy is explained in part by toxic levels of accumulated hydrogen sulfide. Our results demonstrate the existence of an alternative pathway for homocysteine production in *S. cerevisiae*, with implications for a broad range of studies that presume organosulfur auxotrophy in *met15*Δ cells.

## RESULTS

### *met15*Δ cells show unexpected growth in the absence of exogenous organosulfurs

We serendipitously observed robust growth of BY4741 when cells were automatedly transferred in a 384-colony array (**Figure 1A, Table S1**) from a rich medium (YPDA) to a “synthetic defined” (SD) medium lacking the sulfur-containing amino acids methionine (Met) and cysteine (Cys), as well as any other organosulfur, and containing glucose as the carbon source (hereafter referred to as SD-Met-Cys+Glu) (**Figure 1B**). This growth was apparent from the first days of incubation, with colonies reaching saturation after ∼360hrs (**Figure S1**). Both BY4741 and BY4742 (a *MET15*^*+*^ control) harbor complete deletions of *URA3* and *LEU2* (5), rendering them auxotrophic for uracil (Ura) and leucine (Leu). There was a large, statistically significant difference in relative colony size between BY4741 colonies that grew on SD-Met-Cys+Glu and the very small colonies detectable on SD+Met-Cys-Ura+Glu or SD+Met-Cys-Leu+Glu (**Figure 1C**; effect size = 0.627, p-value = 0.021 [Kruskal-Wallis test] and effect size = 0.757, p-value = 0.007 [Kruskal-Wallis test], respectively). In this automated colony transfer procedure, BY4741 does not behave as an organosulfur auxotroph despite lacking *MET15*.

**Figure 1.**
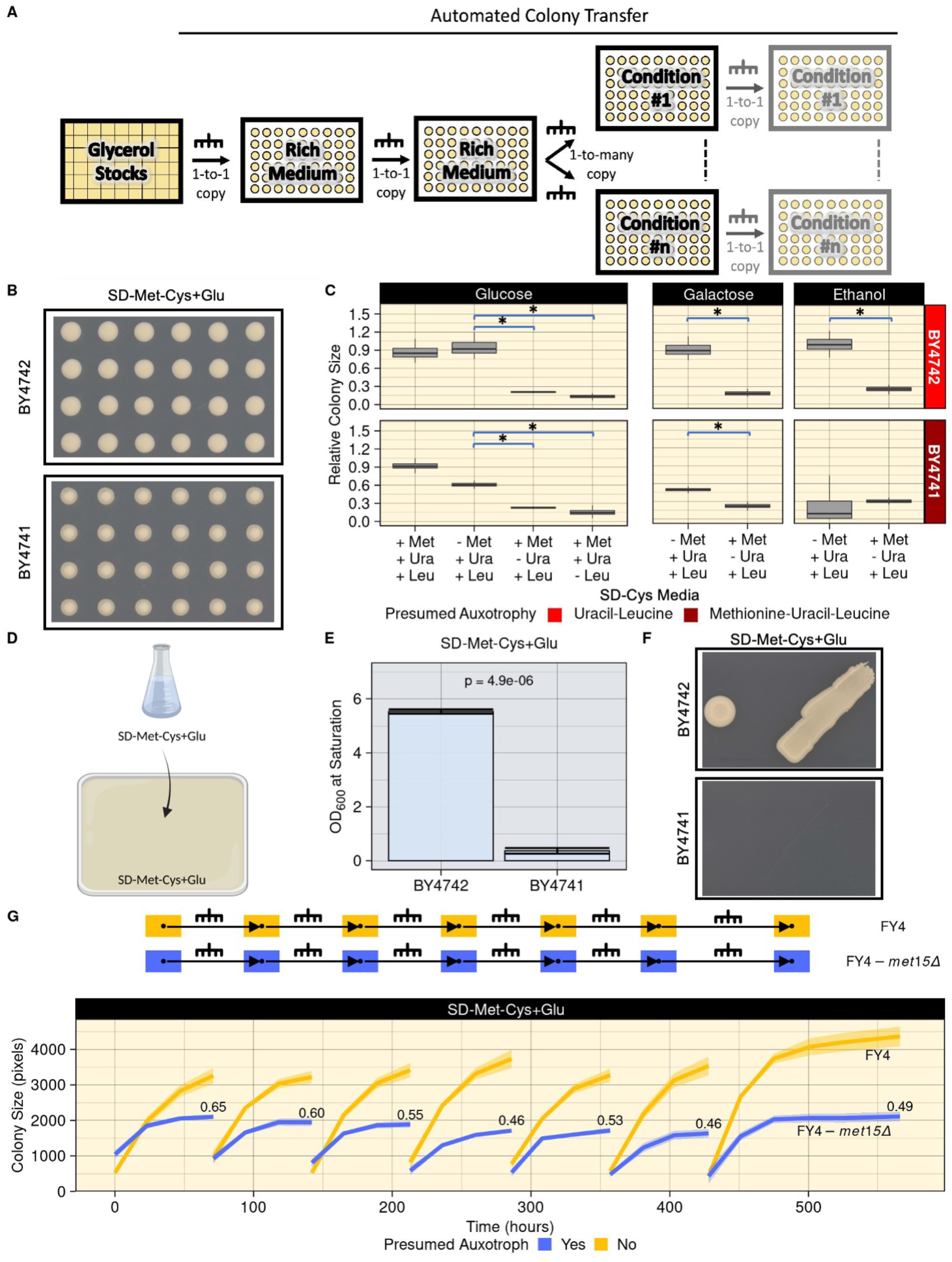
The unexpected growth of *met15*Δ cells on organosulfur-free media is both stable and dependent on cell propagation method. **A)** Automated colony transfer technique used in this study. For some experiments, cells were transferred to the condition plate once; for others, cells were transferred to the condition media multiple times (light gray plates). **B)** Unexpected growth of a strain harboring the *met15*Δ*0* deletion (BY4741) on SD-Met-Cys+Glu. Cropped images of representative 384-density plates after 366 hours of growth are shown. **C)** Robust growth of *met15*Δ*0* cells (BY4741) on SD-Met-Cys+Glu and SD-Met-Cys+Gal, but not on SD-Met-Cys+EtOH. Relative colony size distribution (y-axis) at saturation (∼358 to 366 hours) of BY4742 and BY4741 grown on SD media with glucose, galactose or ethanol, and containing or lacking the indicated nutrients (x-axis). Relative colony size is normalized to the prototrophic FY4 grown in the same conditions (**Figure S3B**). *: significant Kruskal-Wallis test (p-value < 0.05) comparing the median relative colony sizes of biological replicates between two strains. **D)** Experimental setup for traditional cell propagation techniques used in this study. **E)** BY4741 cells fail to grow in liquid SD-Met-Cys+Glu medium. Mean OD_600_ after ∼51 hours in SD-Met-Cys+Glu liquid cultures over three biological replicates for each strain. Error bars: standard error. p: two-tailed Student’s T-test. **F)** BY4741 cells fail to grow on solid SD-Met-Cys+Glu medium in large spots or patches made with a micropipette tip. Cropped representative images of cells grown for 192 hours are shown. **G)** Growth of FY4-*met15*Δ on SD-Met-Cys+Glu is stable over six consecutive automated colony transfers. Top: experimental design. Bottom: colony size (y-axis) over time (x-axis). Lines represent the mean colony size at each time point. Shaded area: one standard deviation. Numbers next to the blue curves indicate the fitness of FY4-*met15*Δ relative to FY4 as determined by dividing median colony sizes at the final time point of each curve.

To confirm this unanticipated observation, we first ensured that our media were devoid of organosulfur contamination using mass spectrometry (**Table S2**). We next performed a cross between BY4741 and BY4742 and dissected nine four-spore tetrads. When patched to SD-Met-Cys+Glu, we observed 2:2 segregation in all tetrads, with two patches growing normally and two patches displaying extremely weak growth, consistent with a monogenic phenotype and indicating that our observations were likely not the result of a secondary mutation (**Figure S2**). The extremely weak growth of patches of cells lacking *MET15* was consistent with auxotrophy, in contrast to the robust growth observed with automated colony transfer. To further investigate the impact of cell propagation technique on growth, we grew BY4741 and BY4742 in liquid SD-Met-Cys+Glu medium and used drops from these cultures to make spots and patches on SD-Met-Cys+Glu solid medium (**Figure 1D**). BY4741 cells grew very little in liquid cultures and not at all in spots or patches, while BY4742 cells grew normally (**Figure 1E-F**). Hence, BY4741 displays auxotroph-like growth when propagated using traditional techniques, but not when automatedly transferred in a grid to seed colonies.

We next interrogated the extent to which the unexpected growth of BY4741 colonies depends on the specific composition of the growth medium. We repeated the automated colony transfer procedure using SD-Met-Cys and SD+Met-Cys-Ura media with either galactose (SD-Met-Cys+Gal, SD+Met-Cys-Ura+Gal) or ethanol (SD-Met-Cys+EtOH, SD+Met-Cys-Ura+EtOH) as the carbon source (**Figure 1C**). As with glucose, when galactose was used as a carbon source, BY4741 colonies grew to much larger sizes on SD-Met-Cys+Gal than on SD+Met-Cys-Ura+Gal (effect size = 0.561, p-value =0.021 [Kruskal-Wallis test]). In contrast, BY4741 displayed miniscule growth on SD-Met-Cys+EtOH. Robust growth was observed in “synthetic complete” (SC) medium containing all amino acids except Met and Cys (SC-Met-Cys) with either glucose or galactose as carbon source (**Figure S3A**).

To study the consequences of *MET15* loss independent of the other auxotrophies present in BY4741, we used CRISPR to make a scarless deletion of *MET15* in the prototrophic FY4 background (FY4-*met15*Δ). We used this strain for all subsequent experiments. Like BY4741, FY4-*met15*Δ cells grew robustly when automatedly transferred to SD-Met-Cys with glucose or galactose, but not ethanol, as carbon sources (**Figure S3B**). FY4-*met15*Δ also grew similar to BY4741 on SC-Met-Cys medium with glucose or galactose (**Figure S3A**). Thus, the unexpected growth of *met15*Δ cells propagated via automated colony transfer is not dependent on the genetic background particular to BY4741.

To determine whether the robust growth of *met15*Δ colonies may be explained by *met15*Δ recycling and/or scavenging of preexisting organosulfurs, we transferred FY4-*met15*Δ from and to SD-Met-Cys+Glu six consecutive times. FY4-*met15*Δ cells displayed robust growth on SD-Met-Cys+Glu over repeated transfers (**Figure 1G**). Relative to FY4, the fitness of these colonies was 0.65 on the initial SD-Met-Cys+Glu plates and stabilized around ∼0.50. The amount of new biomass generated throughout this experiment strongly suggests that the growth of FY4-*met15*Δ cells cannot be explained by recycling or scavenging. Despite a longstanding assumption of auxotrophy, *met15*Δ strains can grow stably in media lacking organosulfurs when propagated via automated colony transfer.

### Stable growth of *met15*Δ cells on organosulfur-free media requires inorganic sulfates and is not seen in other presumed auxotrophs

To investigate whether the unexpected growth phenotype of *met15*Δ strains extends to other key enzymes in the organosulfur biosynthesis pathway, we again used CRISPR to make complete deletions in the FY4 background of nine additional genes (**Figure 2A)**.

**Figure 2.**
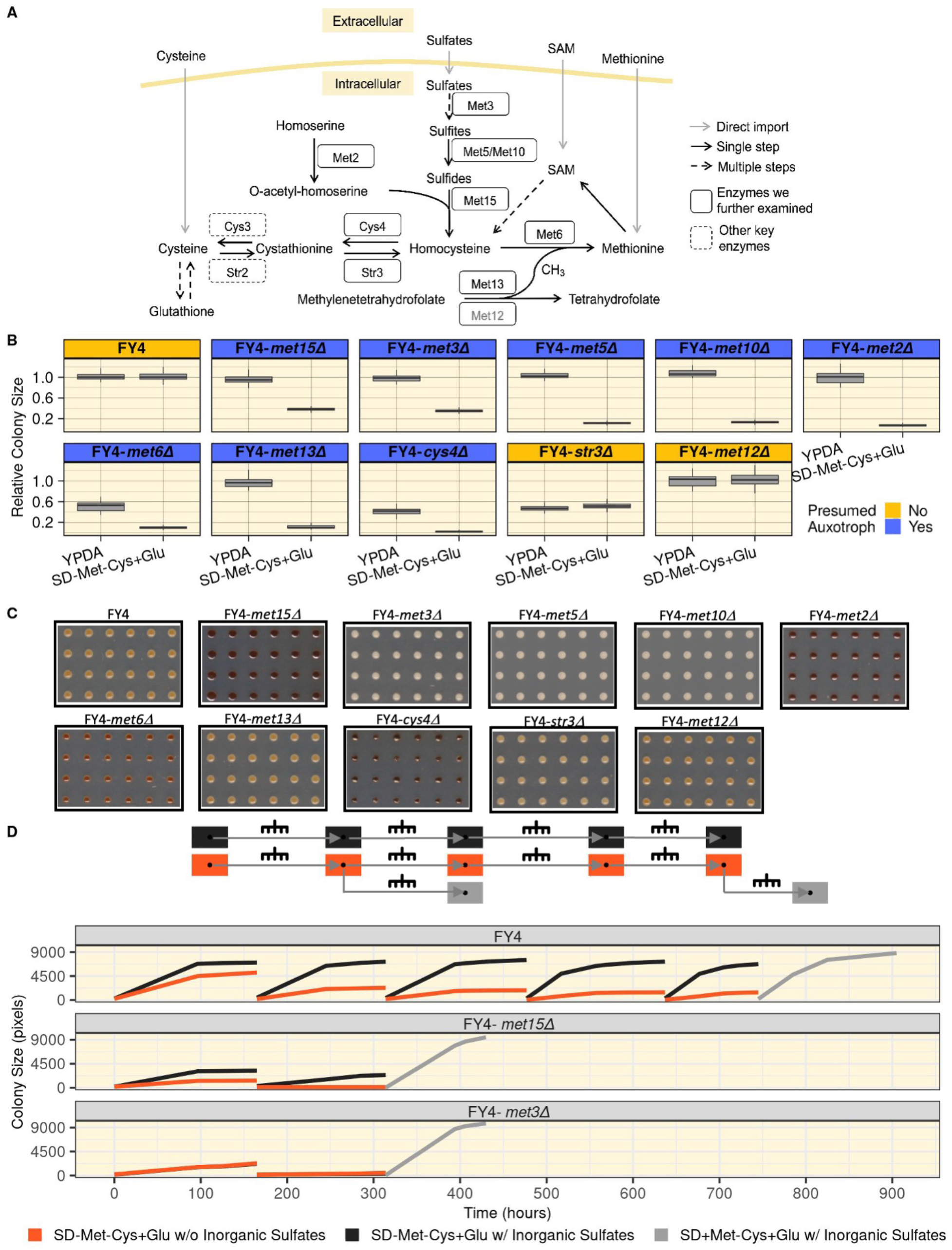
Growth of *met15*Δ colonies in organosulfur-free media is specific and dependent on the utilization of inorganic sulfates. **A)** The *S. cerevisiae* organosulfur biosynthesis pathway. Exogenous organosulfurs such as methionine, cysteine, and SAM can be directly imported (gray arrows); inorganic sulfates can also be imported and reduced to sulfides via the SAP. Sulfides are converted into organosulfurs by Met15, which catalyzes *de novo* homocysteine biosynthesis. Homocysteine is then converted to critical downstream organosulfurs, a selection of which are indicated. *in vivo* methionine biosynthesis requires both the methionine synthase Met6 and the methylenetetrahydrofolate reductase Met13. The Met13 isozyme, Met12 is shown in gray. Enzymes are indicated within boxes, next to the reactions they catalyze, which are illustrated with arrows indicating reaction directionality. Drawing inspired by (52). **B)** *met3*Δ cells also show unexpected growth. Box plots show relative colony size data (y-axis, normalized to FY4) at saturation (∼160-173.5 hours in SD-Met-Cys+Glu; 83.5-89 hours in YPDA). **C)** Disruptions of the organosulfur biosynthesis pathway result in a range of H_2_S accumulation in cells. Representative crops of plate images of strains transferred to complete medium containing bismuth (BiGGY) for ∼160-173.5 hours. **D)** Growth of FY4-*met15*Δ colonies requires inorganic sulfates. Top: experimental design. Bottom: growth curves (colony size, y-axis) of the indicated strains (subpanels) through the entire experimental procedure (time, x-axis) with the mean colony size depicted. The orange (SD-Met-Cys+Glu without inorganic sulfates) and black curves (SD-Met-Cys+Glu with inorganic sulfates) in the lower subpanel (FY4-*met3*Δ) overlap.

These mutants, along with FY4 and the FY4-*met15*Δ strain, were automatedly transferred (**Figure 1A**) to either YPDA or SD-Met-Cys+Glu plates. All mutant strains displayed growth (or lack thereof) on SD-Met-Cys+Glu that was consistent with current knowledge of their roles in organosulfur biosynthesis, with the exception of FY4-*met3*Δ (**Figure 2B**). Met3 catalyzes the first step in the reduction of intracellular sulfate and is thought to be essential for growth when unreduced sulfates are the sole sulfur source (2); however, the FY4-*met3*Δ mutant displayed growth on SD-Met-Cys+Glu that was roughly equivalent to that seen with the FY4-*met15*Δ strain (effect size = 0.651, p-value < 0.001 [Kruskal-Wallis test] and effect size = 0.618, p-value < 0.001 [Kruskal-Wallis test], respectively; **Figure 2B**). The growth of FY4-*met3*Δ was largely unaffected by carbon source (**Figure S4**).

This surprising growth of *met3*Δ cells led us to examine the level of free sulfides present in our mutant strains. To do so we utilized a bismuth assay (13) where the darkness of the colonies is a proxy readout for sulfide accumulation. On a complete bismuth-containing medium (BiGGY), we observed colony colors (**Figures 2C and S5B**) consistent with the known roles of the deleted genes in organosulfur biosynthesis (**Figure 2A**). In particular, *met15*Δ colonies were very dark, consistent with an accumulation of sulfides expected from impaired homocysteine biosynthesis activity, and *met3*Δ colonies were very light, confirming that the SAP is disrupted in these cells as expected. We saw consistent results in a bismuth-containing medium lacking organosulfurs (SD-Met-Cys+Glu+Bi; **Figure S5A**). In this medium, FY4 and FY4-*met15*Δ colonies were darker than in BiGGY (**Figure S5B**), likely resulting from the well-characterized upregulation of the SAP triggered by methionine restriction (8, 14). In contrast, FY4-*met3*Δ remained as light as in BiGGY, confirming again that the SAP is disrupted in these cells as expected (**Figure S5B**). Therefore, the unexpected growth of FY4-*met15*Δ and FY4-*met3*Δ cells is not explained by previous mischaracterizations of the positions of the Met15 and Met3 enzymes in the organosulfur biosynthesis pathway.

To determine whether the surprising growth of FY4-*met15*Δ and FY4-*met3*Δ cells in media lacking organosulfurs depends on the utilization of inorganic sulfates, we examined how these mutants grew over repeated transfers onto plates entirely devoid of both organosulfurs and inorganic sulfates. The growth of FY4-*met15*Δ cells was substantially reduced at first, and essentially abolished upon a second transfer; FY4-*met3*Δ cells failed to show any substantial growth upon a second transfer to medium lacking organosulfurs, irrespective of the presence or absence of inorganic sulfates (**Figure 2D**). Growth was restored for both the FY4-*met15*Δ and FY4-*met3*Δ strains that had stopped growing when we transferred them to SD medium containing both methionine and sulfates (**Figure 2D**). An increase in colony size was also observed for FY4, which had surprisingly continued to grow on media lacking both organosulfurs and inorganic sulfates even after five successive transfers, albeit to a significantly reduced colony size (**Figure 2D**). Therefore these cells, when starved for inorganic sulfates, tend to arrest rather than die, consistent with previous studies (15). Altogether, these results show that deletion of neither Met3 nor any other critical enzyme in the pathway generate the same unexpected growth phenotype as deletion of Met15, and that the unexpected stable growth of *met15*Δ cells requires utilization of inorganic sulfates.

### YLL058W encodes an alternative homocysteine synthase

Our observations led us to hypothesize that the *S. cerevisiae* genome encodes at least one enzyme other than Met15 capable of catalyzing *de novo* homocysteine biosynthesis. We therefore examined the targets of the sole known transcriptional activator of the sulfur metabolic network, Met4 (16). Two of these targets – YHR112C and YLL058W -do not have a known function according to the Saccharomyces Genome Database (SGD) (17) but are predicted by sequence analysis to belong to the same PLP-dependent (pyridoxal 5’-phosphate) transferase superfamily as Met15 (18). Of these two, YLL058W was the more compelling candidate because it has recently been implicated in sulfur metabolism by quantitative trait locus mapping (19) and encodes a protein predicted to have the ability to catalyze homocysteine biosynthesis by a yeast genome-scale metabolic model (20).

YLL058W is located near the telomere of chromosome XII within a cluster of genes that are transcriptionally regulated by Met4, several of which have a direct role in sulfur metabolism (**Figure 3A**) (16). There has been speculation that it may have arisen as a result of horizontal transfer (16) - a relatively rare occurrence in yeast. YLL058W has two close paralogs in the *S. cerevisiae* genome: *STR2*, which encodes an enzyme that converts cysteine into cystathionine (**Figure 2A**), and the uncharacterized YML082W, which is a copy of *STR2* produced during the whole-genome duplication event (21). We generated and clustered a collection of YLL058W, *STR2* and YML082W homologs from bacterial and fungal genomes on the basis of sequence similarity. If YLL058W were derived from a recent horizontal transfer event, we would expect it to be more similar to some of its homologs outside *Ascomycota* than to its *Ascomycota* homologs. Instead, all homologs outside *Ascomycota* grouped together along with all budding yeast homologs more distantly related to *S. cerevisiae* than the *Hanseniaspora* genus. The homologs found in all descendants of the common ancestor of *Saccharomyces* and *Hanseniaspora* comprised two additional groups, one containing YLL058W and the other containing both *STR2* and YML082W (**Figure S6**). The genes belonging to the YLL058W class and the *STR2*/YML082W class appear at approximately the same point in the phylogenetic tree (**Figure 3B**), prior to whole-genome duplication. Therefore, we infer that an ancestral gene duplicated into the YLL058W and *STR2/*YML082W ancestors, followed by substantial sequence divergence of both copies, and later by duplication of the *STR2/*YML082W ancestor. Interestingly, in all species that contain genes of the YLL058W class, the YLL058W homolog is located near the telomeres among a collection of other genes related to sulfur metabolism (**Figure 3A-B and Table S3**). The preservation of this pattern in the genomically-unstable near-telomere region is consistent with a strong selective constraint keeping YLL058W in proximity to other genes involved in sulfur metabolism.

**Figure 3.**
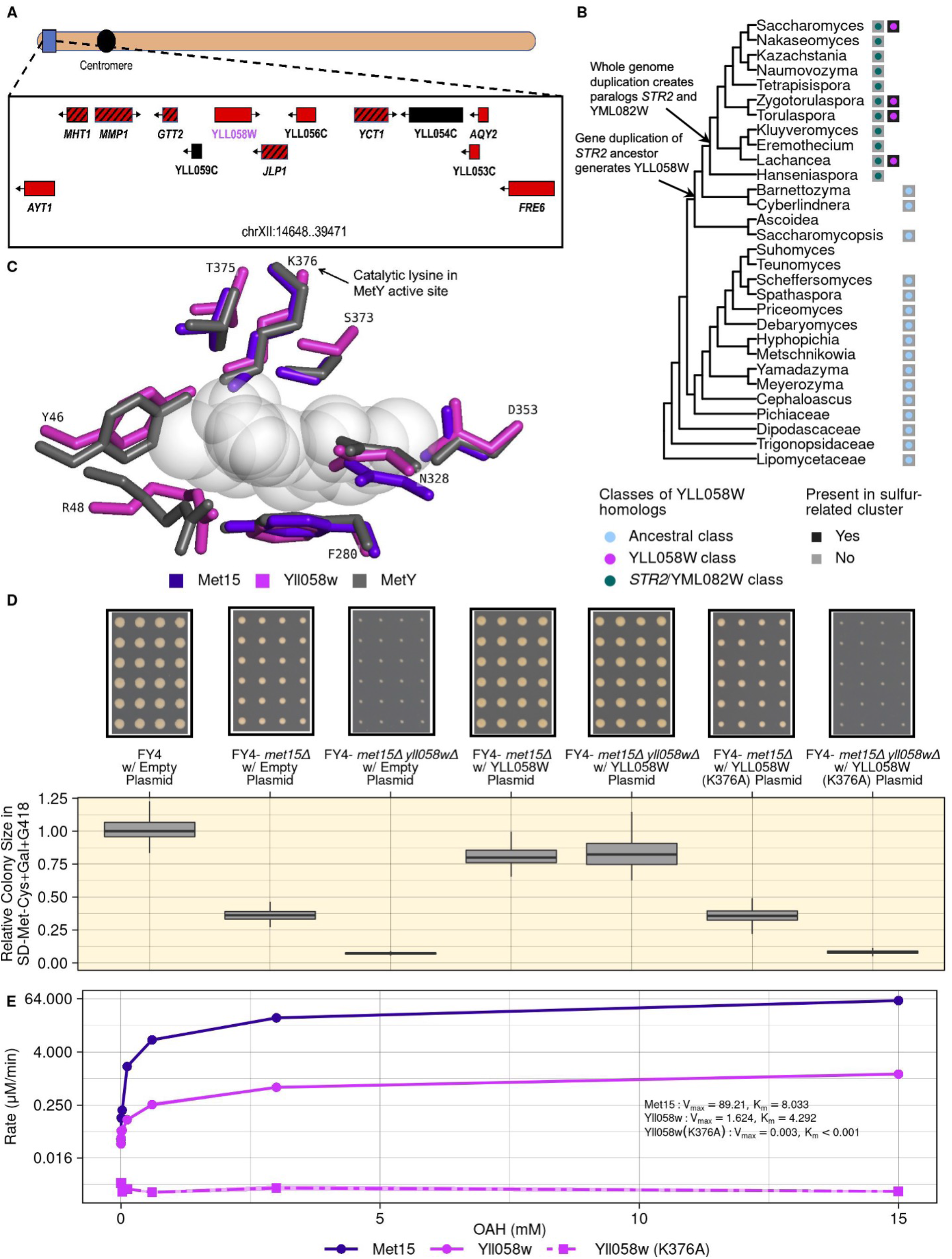
Yll058w functions as an inefficient homocysteine synthase. **A)** Genomic context of YLL058W (17). In S. cerevisiae, YLL058W (magenta) is located near a cluster of genes that are transcriptionally activated by Met4 ((16) red boxes), several of which have a direct role in sulfur metabolism (red boxes with black stripes). Black boxes indicate ORFs that fall into neither category. **B)** Distribution of YLL058W homolog clusters across the budding yeast phylogenetic tree. The tree topology follows (47). Circles adjacent to each taxon indicate whether YLL058W homologs belonging to each of three classes are found in the genome of that taxa, while black boxes indicate the presence of a cluster of sulfur-related genes, as depicted in Panel A for *S. cerevisiae*. **C)** Yll058w has structural similarity to a homocysteine synthase. Overlap of the active site residues of MetY from Thermatoga maritima (PDB ID: 7KB1) (gray) with the predicted structures of Yll058w (magenta) and Met15 (violet), calculated using AlphaFold, are shown. A reaction intermediate captured within the MetY structure (23) is shown in semitransparent spheres. Residue labeling corresponds to the Yll058w sequence. **D)** Yll058w is critical for growth in the absence of *MET15*. The indicated strains bearing high-copy (2μ ORI) plasmids either empty or overexpressing YLL058W or YLL058W (K376A), were grown on SD- Met-Cys+Gal+G418 medium. The box plots show relative colony sizes at saturation (296 hours). Cropped, representative images of the plates corresponding to each strain are shown above. **E)** Yll058w is a less efficient homocysteine synthase. Enzyme kinetic assay was carried out using the recombinantly expressed purified proteins used in **Figure S9**, Met15 purple solid line, Yll058w pink solid line and the putative catalytic dead mutant of Yll058w (pink dashed line). Varying concentrations of the substrate, O-acetyl Homoserine (OAH), are shown on the x-axis and the rate of reaction on the y-axis. Inferred V_max_ and K_m_ values of Met15, Yll058w and Yll058w (K376A) are shown in the panel and **Supplementary Table S10**.

We next examined the predicted structure of the putative Yll058w enzyme. To this aim, we compared structures predicted by AlphaFold2 (22) for Yll058w and Met15 to the experimentally determined crystal structure of MetY (**Figure S7A**), a bacterial Met15 ortholog that catalyzes homocysteine biosynthesis from OAH and bisulfide (23). The MetY crystal structure captured a reaction intermediate, allowing for a precise determination of active site residues. The predicted structures of both Met15 and Yll058w are highly similar to MetY within the active site, including the catalytic lysine K376 (**Figures 3C and S7B**). MetY and Met15 function as homotetramers (23, 24), with multimerization domains allowing the distinct monomers to come together and complete the active site (**Figure S7C**). In contrast, Yll058w lacks a multimerization domain and is predicted to be monomeric. However, Yll058w contains an approximately 180 amino acid region that has no homology to MetY or Met15. This region is predicted to fold back onto the core of the protein, supplying the missing amino acids to complete the active site (**Figure S7D**). Altogether, these comparative structural analyses suggest that a Yll058w monomer may be capable of carrying out catalysis similar to that of the MetY and Met15 tetramers.

We next employed a genetic approach to determine whether the putative Yll058w enzyme is necessary and sufficient to support the growth of *met15*Δ cells in the absence of exogenous organosulfurs. We used CRISPR to delete YLL058W and to make a *met15*Δ *yll058w*Δ double mutant in the FY4 background. We also constructed two high copy-number plasmids, one overexpressing YLL058W, and the other overexpressing a YLL058W mutant in which the putative catalytic lysine (K376, **Figure 3C**) was changed to an alanine (A376). The FY4-*yll058w*Δ single mutant resulted in only a very modest growth defect on SD-Met-Cys+Glu (effect size = 0.054, p-value > 0.05 [Kruskal-Wallis test]) (**Figure S8A**), and wildtype levels of H_2_S accumulation (**Figure S8B**). Strikingly however, the FY4-*met15*Δ *yll058w*Δ double mutant showed little to no growth in the absence of exogenous organosulfurs (**Figure 3D**), indicating a synthetic lethal interaction between *MET15* and YLL058W. Overexpression of YLL058W fully compensated for the loss of *MET15*, both alone (effect size= 1.238, p-value < 0.001 [Kruskal-Wallis test]) and in combination with the loss of YLL058W (**Figure 3D**; effect size= 10.117, p-value < 0.001 [Kruskal-Wallis test]). In contrast, the overexpression of YLL058W (K376A) in these same strain background has no rescue effect (**Figure 3D**). These results strongly suggest that the growth of *met15*Δ cells in the absence of exogenous organosulfurs is mediated by the enzyme encoded by YLL058W.

To directly examine the ability of Yll058w to catalyze homocysteine biosynthesis, we first purified recombinant Yll058w and Yll058w (K376A) proteins from *E. coli*, along with Met15 as a positive control (**Figure S9**). We then carried out *in vitro* reactions using the canonical O-acetyl-homoserine (OAH) and H_2_S substrates as well as PLP, a cofactor critical for Met15 activity. Yll058w was able to catalyze homocysteine production significantly above background, albeit much less efficiently than Met15, with the Met15 reactions yielding ∼9.9 times as much homocysteine at the 15-minute time point (**Figure S10A**). We then measured enzyme kinetic parameters by performing analogous reactions with varying amounts of the limiting substrate, OAH (**Figure 3E**). Yll058w activity was consistently above background and increased with increasing OAH concentrations (V_max_: 1.62; K_m_: 4.29), although remaining less efficient than Met15 activity at all OAH concentrations (V_max_: 89.21; K_m_: 8.03). The putative catalytic dead mutant Yll058w (K376A) had no detectable activity. We therefore conclude that YLL058W encodes an inefficient homocysteine synthase capable of catalyzing the same reaction as Met15.

To investigate whether the inefficient enzymatic activity of Yll058w would be sufficient to support growth of *met15*Δ cells in the absence of exogenous organosulfurs, we adapted an enzyme-constrained version of the consensus genome-scale metabolic model for *S. cerevisiae*, Yeast8 (25). We removed *MET15* from the model, added the Yll058w reaction we observed *in vitro*, and evaluated the flux through the biomass equation at a range of arbitrarily assigned catalytic efficiencies (Kcat) for Yll058w. We found that an appreciable reduction in biomass occurred only when Yll058w was assigned a catalytic efficiency ∼100x lesser than that assigned to Met15 in the model (**Figure S10B**). These simulations, together with our structural, *in vivo*, and biochemical findings, demonstrate that it is plausible that Yll058w supports growth of *met15*Δ cells in the absence of exogenous organosulfurs.

### Removal of H_2_S enhances growth of *met15*Δ cells in organosulfur-free media

We hypothesized that the large excess of H_2_S present in *met15*Δ cells (**Figures 2C and S5**) may explain why these cells do not grow without organosulfurs using traditional growth techniques despite having the capacity to synthesize homocysteine through Yll058w. This accumulation of H_2_S is expected since the reaction catalyzed by Met15 is one of the major sulfide-consuming reactions in the cell (26), and since methionine restriction triggers upregulation of the SAP (8, 14). Although some amount of H_2_S is required for cell growth, at high levels it is toxic in a range of species (27-29), including in *S. cerevisiae* (30). To investigate this hypothesis, we examined the growth of FY4 and FY4-*met15*Δ cells in liquid SD-Met-Cys+Glu cultures with or without the H_2_S chelator ferric (Fe)-EDTA, which bonds free sulfides that exit the cell by diffusion (31, 32) (**Figure 4A**). In the presence of Fe-EDTA, we observed a modest increase in the growth of FY4 cultures (p-value = 0.041 [Student’s t-test]; **Figure 4B**) and a striking increase in the growth of FY4-*met15*Δ cultures (p-value = 0.01 [Student’s t-test]; **Figure 4B**).

**Figure 4.**
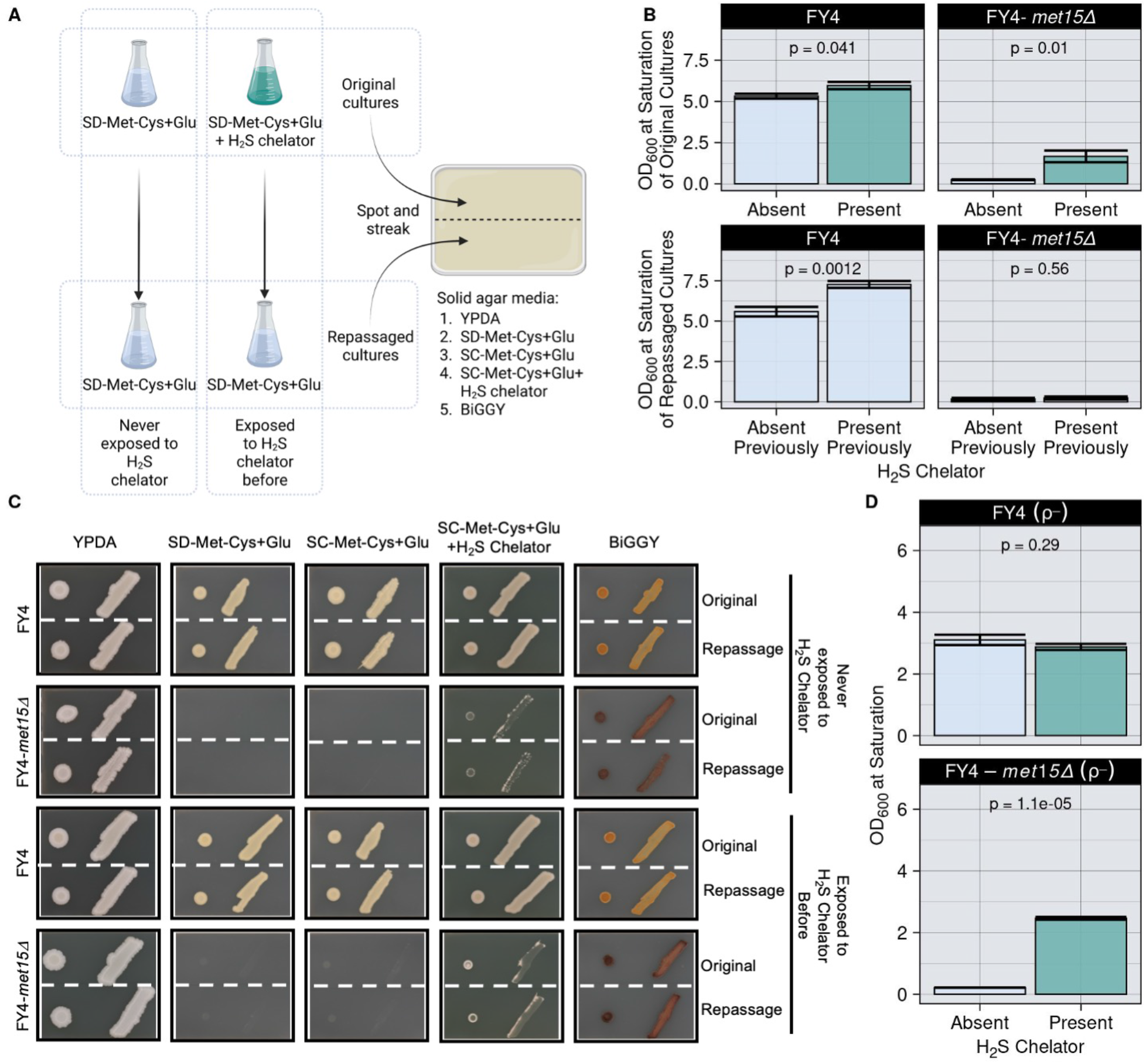
Chelation of H_2_S facilitates the growth of *met15*Δ cells in the absence of exogenous organosulfurs. **A)** Experimental setup. Strains were grown in either SD-Met-Cys+Glu or SD-Met-Cys+Glu+Fe-EDTA liquid media (top two flasks, original cultures). Once saturation was reached for all cultures, cells were repassaged in SD-Met-Cys+Glu media (bottom two flasks, repassaged cultures) and again grown to saturation. Ten μl from each culture were then spotted and struck to the indicated solid media using a micropipette tip. **B)**The H_2_S chelator Fe-EDTA partially rescues growth of *met15*Δ cells in liquid media lacking organosulfurs. Mean OD_600_ values of the cultures after ∼51 hours of growth with or without chelator (top panel). Exposure to the chelator does not induce heritable growth-enabling mutations (lower panel). Mean OD_600_ values of the cultures after ∼51 hours of growth after repassage. Error bars: standard error; p: two-tailed, unpaired Student’s t-test. Individual values for each replicate, from both independent experiments, are shown in **Table S4**; outlier FY4-*met15*Δ cultures that showed unexpected growth in SD-Met-Cys+Glu were excluded from the data presented here and indicated in green text in the table. **C)** Chelation of H_2_S partially rescues the growth of FY4-*met15*Δ cells grown in spots or patches (see **Panel A**). Cropped, representative images of cells grown for 192 hours are shown. **D**) Growth defect of *met15*Δ petite (ρ-) derivatives in liquid media lacking organosulfurs (SD-Met-Cys+Glu) is rescued in presence of H_2_S chelator. Data generation was performed as in the original cultures from **Panel A**, data representation is same as in **Panel B**.

To rule out spontaneous mutations as the reason for the growth increase of FY4-*met15*Δ cultures, we repassaged the cultures in SD-Met-Cys+Glu medium lacking chelator (**Figure 4A**). The repassaged FY4-*met15*Δ cultures showed little to no growth (**Figure 4B**). Interestingly, the repassaged FY4 cultures that had been propagated in the presence of chelator saturated at a higher OD_600_ than those that had not (p-value = 0.0012 [Student’s t-test]; **Figure 4B**). We then placed cells from both the original and the repassaged cultures on solid media plates in large drops and patches (**Figure 4C**). The FY4-*met15*Δ cells failed to grow on SD-Met-Cys+Glu and SC-Met-Cys+Glu in drops or patches, irrespective of their previous exposure to the chelator (**Figure 4C**). These results demonstrate that the growth of FY4-*met15*Δ cultures in the presence of chelator is not facilitated by heritable mutations. Growth in drops and patches of FY4-*met15*Δ cells was partially rescued by adding Fe-EDTA to the organosulfur-free medium (**Figure 4C**). Thus, the growth defect (in some settings) and auxotrophic-like behavior (in other settings) of *met15*Δ cells grown without organosulfurs is at least partially attributable to H_2_S toxicity.

Excess H_2_S is often associated with mitochondrial dysfunction (33). Yeast strains with damaged or absent mitochondrial genomes display the so-called “petite” phenotype (34). To determine whether the partial rescue of *met15*Δ cells by an H_2_S chelator was dependent on the mitochondrial genome, we constructed petite (ρ-) derivatives of FY4 and FY4*-met15*Δ and grew them in SD-Met-Cys+Glu cultures with or without Fe-EDTA. We observed a strong rescue of FY4-*met15*Δ ρ-growth upon addition of the chelator (p-value = 1.1 ×10^−5^ [Student’s t-test]; **Figure 4D**, bottom panel). This result indicates that the chelator-mediated rescue of *met15*Δ cells growth in liquid cultures does not require an intact mitochondrial genome. Next, we examined the growth of these petite strains and their respective non-petite parental strains in our automated colony transfer pipeline (**Figure 1A**). The cells were transferred to solid media containing or lacking methionine, and with either glucose or ethanol as the carbon source (**Figure S11**). In SD+Met-Cys+EtOH, non-petite FY4-*met15*Δ cells grew poorly relative to their *MET15*^*+*^ counterparts. In SD-Met-Cys+Glu, the FY4-*met15*Δ ρ-strain grew almost as well as their non-petite parental strain. These results suggest that *met15*Δ cells experience some degree of mitochondrial dysfunction that impairs their ability to metabolize ethanol, but that neither their unexpected growth in automated colony transfer settings, nor the chelator-mediated rescue of their lack of growth in traditional propagation techniques, require a functional mitochondrial genome.

## DISCUSSION

Here we show that, contrary to longstanding assumptions, the organosulfur auxotrophy-like phenotype in a *met15*Δ background is dependent on cell propagation technique. Our analyses demonstrate that the *S. cerevisiae* genome, and possibly those of several closely related fungal species, encode another enzyme, the previously uncharacterized Yll058w, that can catalyze the same reaction as Met15, albeit less efficiently. This activity compensates for the loss of *MET15* and enables the growth of *met15*Δ cells in the absence of exogenous organosulfurs when cells are propagated using automated colony transfer or grown in the presence of an H_2_S chelator. Altogether, our observations support a model according to which *met15*Δ cells can synthesize enough homocysteine to support growth through the activity of Yll058w, but often experience chronic H_2_S toxicity that prevents growth when standard laboratory techniques are used (**Figure 5**). Given its evolutionary conservation, Yll058w may well have other activities that are not investigated here. For example, YLL058W was recently implicated in *S*-methylmethionine metabolism by quantitative trait locus mapping (19).

**Figure 5.**
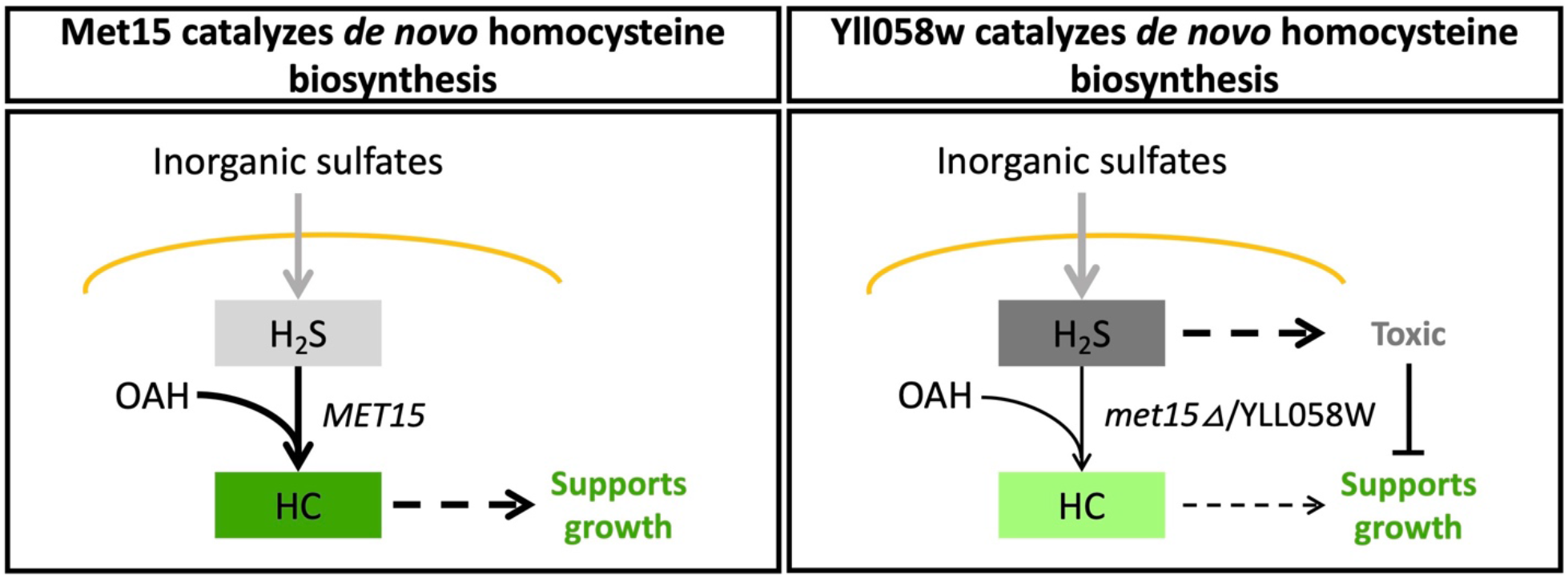
H_2_S toxicity inhibits growth when Yll058w catalyzes *de novo* homocysteine biosynthesis. Model representing the hypothesized response to organosulfur starvation in cells containing or lacking Met15. Extracellular inorganic sulfates are imported through the cell membrane (yellow line) and reduced to H_2_S. When Met15 is present (left), cells efficiently convert H_2_S and O-acetylhomoserine into homocysteine, leading to a moderate amount of H_2_S (light gray) and an adequate amount of homocysteine (HC; dark green) to support normal growth. When Met15 is absent, homocysteine biosynthesis is catalyzed by the comparatively inefficient Yll058w enzyme (right). The resulting lower amount of homocysteine (light green) is sufficient to support growth in some contexts, but the high amount of H_2_S that accumulates (dark gray), exacerbated by an upregulation of the SAP (8, 14), can become toxic and lead to cell death or arrest in other contexts.

### H_2_S-mediated dependence of a classic auxotrophy

H_2_S levels are extremely high when *met15*Δ cells are growing without exogenous organosulfurs (**Figures 2C** and **S5**), as a result of the upregulation of the SAP (8, 14) and Yll058w being an inefficient homocysteine synthase. Our results indicate that H_2_S toxicity is at least partially responsible for the arrest and/or death that occurs in *met15*Δ cells grown using traditional techniques (**Figure 5**). A primary mechanism for the toxic effects of elevated H_2_S, seen in many organisms, involves mitochondrial dysfunction (33). The failure of *met15*Δ cells to utilize ethanol (**Figures 1C, S3B**, and **S11**) is consistent with mitochondrial dysfunction. However, the ability of the H_2_S chelator Fe-EDTA to restore the growth of FY4-*met15*Δ ρ-cells grown in liquid cultures (**Figure 4D**), as well as the ability of these cells to grow when automatedly transferred to solid SD-Met-Cys+Glu medium (**Figure S11**), suggests that H_2_S is limiting the growth of *met15*Δ cells in part through a pathway that does not involve the mitochondrial genome. The precise molecular mechanisms at play may be dissected in future studies. Our discovery that automated colony transfer allows for growth of *met15*Δ cells without organosulfurs provides a new tool to investigate this question as well as, more generally, the response to extreme sulfur starvation and associated phenotypes such as aging.

We speculate that, when cells are grown via automated colony transfer, excess sulfides can be eliminated through diffusion between the evenly spaced small colonies whereas, in thick patches of cells, excess sulfides are secreted and affect neighboring cells, creating group-level toxicity and self-limiting growth. Similarly, we speculate that H_2_S toxicity is high in liquid cultures because the semi-sealed environment limits gas exchange. Interestingly, a recent preprint suggested that when *met15*Δ cells are grown to a high density in liquid cultures, they can keep growing after having exhausted exogenous organosulfurs by utilizing the excess H_2_S secreted in the prior phase of the culture (35). This preprint implicated Yll058w as the key enzyme allowing cells to re-incorporate the sulfide ions and showed that the equilibrium of H_2_S levels between the aqueous (retained in the media and accessible) and gas phases (escaped and inaccessible) determined whether organosulfur auxotrophy could be rescued. Our combined observations hint at a cell-density dependent rescue of organosulfur auxotrophy that is influenced by the interplay between the amount of H_2_S that is: 1) produced by the cells, 2) retained in the microenvironment, and 3) released into the atmosphere. Differences in cell propagation techniques likely introduce unappreciated differences in the dynamics of gas exchange that determine whether *met15*Δ cells grow, arrest or die. While the conditional essentiality of genes has been well studied in the context of differing genetic backgrounds or environments (36, 37), to our knowledge, *MET15* represents the first report of a case in which the essential nature of a gene is dependent on the way cells are propagated.

### An expanded yeast sulfur metabolic network

It is of note that even the prototrophic FY4 strain displayed enhanced growth in liquid media when cells were previously exposed to an H_2_S chelator (**Figure 4B**). While it is possible that Fe-EDTA enhances the growth of even *MET15*^*+*^ strains due to some secondary effect unrelated to the chelation of H_2_S, previous studies showed that very little of the chelator is taken up by yeast cells (31). Furthermore, our data indicate that wild-type strains accumulate H_2_S to a level that is roughly intermediate between that seen in *met15*Δ cells and mutants lacking an intact SAP (**Figures 2C and S5**). This suggests that the possibility that 1) even wild-type cells suffer some degree of H_2_S-induced stress when grown in organosulfur-free media, and 2) removal of excess H_2_S allows the cells to adapt to this stress, likely at the level of gene expression or epigenetics. This may have implications for anyone seeking to maximize cellular growth in media lacking, or containing low amounts of, organosulfurs, whether in an industrial or a research setting. Also surprising was the limited but continued growth of FY4 cells even after multiple successive transfers to media lacking both organosulfurs and inorganic sulfates (**Figure 2D**). This continued growth was unanticipated given that to our knowledge the SAP relies on inorganic sulfates (**Figure 2A**). Perhaps Met15, but not Yll058w, can metabolize derivatives of other sulfur-containing compounds present in our growth media (*e*.*g*. biotin). Altogether, our work suggests that there may yet be more unknown factors that contribute to the yeast sulfur metabolic network.

Sulfur-containing amino acids act as cellular sensors for growth control (38, 39). In particular, methionine restriction is known to increase longevity in a broad range of organisms (40), and may serve as a therapeutic treatment for certain cancers (41). Mutants lacking *MET15* have been widely used in the study of lifespan extension in yeast (8-10, 42). Our data add a potential layer of nuance to such studies, including those that have explored the role for H_2_S specifically in lifespan extension (e.g. (43)). More generally, our discovery of the alternative homocysteine synthase Yll058w may influence the interpretation of the plethora of published studies, including genome-wide screens conducted in the BY4741 background, wherein the results or methodologies assumed *met15*Δ cells to be auxotrophic for organosulfurs. Our findings that *S. cerevisiae* can grow in the absence of exogenous organosulfurs despite very low levels of *de novo* homocysteine biosynthesis, that the previously observed auxotroph-like phenotypes are dependent on cell propagation techniques, and that H_2_S levels control group-level growth behaviors, provide an additional avenue for studying the interplay between sulfur metabolism and its many dependent phenotypes.

## EXPERIMENTAL PROCEDUES

This article contains supporting information, including **Supplementary Figures S1-12** and **Supplementary Tables S1-S10, Supplementary File 1** and **Supplementary Materials and Methods**.

### Yeast strains and growth media

See **Supplementary Materials and Methods** and **Tables S6, S7** and **S8** for a list and description of all strains, plasmids, guide RNAs and repair fragments used for this study. See **Table S9** for recipes for the various growth media used in this study. For each strain we selected four independent colonies from streakouts, which were used to inoculate liquid YPDA medium and grown overnight at 30°C. These cultures were considered biological replicates. The cultures were then used to create 384-well glycerol stocks (glycerol 25% v/v), with each biological replicate occupying one-fourth of the plate, distributed uniformly.

### Automated Colony Transfer

Using the benchtop RoToR HDA robotic plate handler (Singer Instruments Co Ltd, Roadwater, UK) strains were transferred from glycerol stocks to YPDA agar plates and incubated at 30°C for 72 hours (see **Table S1** for a description of the settings used). Colonies on each plate were transferred to two YPDA plates (representing technical replicates) and again incubated at 30°C for 48 hours; this step reduces the variability in colony sizes introduced when cells are transferred from glycerol stocks. Finally, these plates were “one-to-many” transferred onto various solid “condition” media plates. Condition plates were incubated at 30°C and periodically imaged until saturation. This protocol, depicted in **Figure 1A**, was carried out at least twice per experiment to control for batch effect. **Figures 1B-C, 1G, 2B, 2C, 2D, 3D, S1, S3, S4, S5, S8 and S11** were carried out in this manner.

### Relative fitness measurement

A spatially cognizant colony size database was built using colony size estimations from the plate images per the LI Detector framework for high-throughput colony-based screening analysis (44). For all experiments, border colonies were removed from the dataset as these tend to grow more due to greater nutrient access, leaving a maximum of 77 colony size observations per strain per time point per plate per biological replicate. Outlier colony size values were then removed using two median adjusted deviations as a cutoff. These raw colony size values are reported in **Figures 1G** and **2D**. Otherwise, the median colony size per time point of the prototrophic strain FY4 was used to calculate relative colony size of every colony per strain per time point. At saturation, the relative colony size distribution per strain is determined by the median relative colony sizes per plate per biological replicate. This relative colony size distribution is compared amongst strains using the non-parametric Kruskal-Wallis Test to determine significant changes in fitness. Effect size values are calculated as the difference in median colony sizes divided by the median colony size of the strain being used as the reference. Additionally for **Figure S11**, growth curve analysis was done with the *growthcurver* (45) package in R (46), using the time series colony size data per colony per strain per technical replicate per biological replicate. These data were first converted into an equally spaced time series using a second degree locally weighed scatterplot smoothing (loess) function to generate 100 equally spaced data points. This was then fed into the ‘*SummarizeGrowthByPlate()*’ function of the gro*wthcurver* package (45) to estimate the area under the curve (AUC) per colony per strain in the experiment. AUC summarizes a growth curve by integrating the contributions of the initial population size, growth rate, and carrying capacity into a single value, making it a more robust estimation of fitness. Relative AUCs, effect sizes and significant changes were estimated the same way as the colony size data described above. The AUCs from the data in **Figure S11** were used to calculate the statistics in **Table S5**.

### Evolutionary origins of YLL058W

Genome sequences of 332 budding yeasts were taken from a published dataset (47). Each genome was scanned for ORFs (consisting of the sequence between an ATG start codon and a stop codon in the same frame) larger than 300 bp. BLASTP was then run with default settings and a 10^−4^ e-value cutoff on the predicted protein products of these ORFs to identify the strongest match among *S. cerevisiae* annotated genes. All ORFs for which the best match (i.e., the lowest BLASTP e-value) was either the *S. cerevisiae* YLL058W sequence or the sequence of one of its two *S. cerevisiae* homologs (*STR2* and YML082W) were retained. For each homolog identified, we also noted the best *S. cerevisiae* matches for the 5 nearest ORFs in each direction that had a match. To supplement this list with homologs from more distant species, we submitted the *S. cerevisiae* YLL058W sequence to the HMMER homology search tool (48). From these results, we selected the strongest matching sequences (lowest e-values) from a variety of bacterial and fungal taxa outside of the *Ascomycota* phylum (four bacterial and nine fungal outgroups). The full list of YLL058W homologs and their sequences is given in **Supplementary File 1**. We next constructed a multiple alignment of all homologs using the MUSCLE aligner (49). We used the R package bio2mds to perform K-means clustering on the sequence alignment and to visualize the results using multidimensional scaling (**Figure S6**).

### *in vitro* homocysteine assay

*in vitro* homocysteine biosynthesis was carried out largely as described previously (3). Aliquots of Met15, Yll058w and Yll058w (K376A), purified as described (**Supplementary Materials and Methods, Figure S9**), were dialyzed into a final storage buffer of 20 mM Tris-Cl pH 8.0, 500 mM NaCl, 20 mM Imidazole, 0.2 mM pyridoxal 5’-phosphate (PLP), and 10% glycerol, snap frozen, and stored at -80°C. Aliquots were thawed on ice, and the enzymes were diluted in the storage buffer to a concentration of 60 nM and preincubated at room temperature for 10 minutes. To compare enzymatic activity of Met15 with Yll058w at a saturating substrate concentration, the enzyme solution was then mixed with diluted 10x, giving a final enzyme concentration of 6nM, with a reaction mixture containing 50 mM Potassium phosphate buffer (pH 7.8), 1.0 mM EDTA pH 8.0, 15.0 mM O-acetyl-L-homoserine, 5.0 mM NaHS, and 0.6 mM PLP. The reaction components were then mixed by gentle pipetting and incubated at 30°C. Each reaction was performed in triplicate. Samples were removed at 6 timepoints (0, 1, 3, 5, 10, and 15 minutes), and the reactions were stopped by adding 1.0 M HCl at a ratio of 10mL for every 100mL of sample, then snap frozen. To calculate K_m_ and V_max_ for Met15, Yll058w and Yll058w (K376A), the reaction was performed for 10 minutes with a range of O-acetyl-L-homoserine from 0-15mM as indicated. In the same buffer as the reaction mixture but without any enzyme or substrate (but with acid added), we also generated a homocysteine standard curve with seven serial 1:5 dilutions ranging from 15 mM to 0.00096 mM, as well as a 0 mM “blank” sample. Homocysteine levels were measured as described in **Supplementary Materials and Methods**.

### Metabolic Modeling

Metabolic modeling simulations were performed with a Python script making use of the COBRApy toolkit (50) and the ecYeastGEM_batch model (25), an “enzyme-constrained” variant (51) of Yeast8. The enzyme-constrained feature of the steady-state model is achieved by treating enzymes as metabolites that are temporarily “consumed” (i.e. removed from the available pool) while they are engaged in catalysis. The script first removes all preexisting reactions (which have not been experimentally validated) contributing to the “consumption” of Yll058w (r_0815No1, r_0815REVNo1, and arm_r_0815). It then adds the homocysteine-generating reaction that we observe in our *in vitro* assay, which uses H_2_S and *O*-acetylhomoserine as substrates and yields homocysteine, acetate, and hydrogen as products, with Yll058w as the catalyst. This reaction was added with a range of Kcat values assigned to Yll058w in order to evaluate the “biomass” generated over a broad range of enzyme efficiencies. The reaction catalyzed by Yll058w was made to be bidirectional; however, flux was only observed in the forward (homocysteine-producing) reaction. *MET15* was then removed from the model, eliminating all reactions that the encoded enzyme was associated with. Finally, the model was “solved” to obtain the flux through the biomass equation, a proxy for fitness. The default growth medium associated with the model, which contains sulfates but lacks organosulfurs, was used for all simulations.

### Chelation of H_2_S with Ferric-EDTA sodium salt

For each strain, five single colonies were scraped from a YPDA agar plate and cells were resuspended in SD-Met-Cys+Glu medium. Optical density at 600 nm (OD_600_) of the suspension was measured, and 25mL cultures were inoculated at a starting OD_600_ of 0.1 in SD-Met-Cys+Glu or SD-Met-Cys+Glu supplemented with 0.048M Ferric (Fe)-EDTA (Ethylenediaminetetraacetic acid ferric sodium salt, E6067, Millipore Sigma) in 125mL Erlenmeyer flasks (optimal Fe-EDTA dosage was determined by a titration experiment (**Figure S12**)). For the non-petite strains, the experiment was performed twice; each time, cultures were prepared in triplicate. The petite strains were tested in a single experiment using four biological replicates. Cultures were incubated at 30°C with 150rpm shaking and OD_600_ was measured regularly until saturation was reached (∼51 to 53 hours). For the repassaging of the non-petite strains (**Figures 4A** and **4B-C**), cells were then pelleted to facilitate the exchange to fresh SD-Met-Cys+Glu medium, again using a starting OD_600_ of 0.1, and incubated as before for ∼51 hours with regular OD_600_ measurements. A portion of each culture was stored at 4°C. Ten μl drops of each original and repassaged culture were then both spotted with a pipette, and struck with a sterile pipette tip, on agar plates (YPDA, SD-Met-Cys+Glu, SC-Met-Cys+Glu, SC-Met-Cys+Glu+(Fe-EDTA) and BiGGY) that were incubated at 30°C (**Figure 4C**) and imaged regularly with spImager Automated Imaging System (S & P Robotics Inc., Ontario, Canada).

## DATA AVAILABILITY

All data generated/analyzed in this study are available in the main text, in the Supplementary Figures and Tables, and as Supplementary Data files. Supplementary material is available at https://bit.ly/3GIgpld. All images and statistical analyses in this study were generated using code available at: https://github.com/sauriiiin/methionine/tree/master/paper.

## ACKNOWLEDGEMENTS

We thank all the members of the Carvunis Lab, and in particular Carly Houghton, for helpful suggestions during lab meetings and presentations. We also thank Karen Arndt for helpful discussions, Kate Metedgul-Ernar, Cameron Hines and Brian Hsu for their precious help in the early stages of the project and S.G. Wendell and Steven Mullett for their assistance with mass spectrometry analysis. We also thank the Moffitt Proteomics/Metabolomics Core, which is funded in part by Moffitt’s Cancer Center Support Grant (NCI, P30-CA076292).

## AUTHOR CONTRIBUTIONS

Conceptualization: S.B.V.O., S.B.P., N.C.C., G.M.D., A-R.C. Methodology: S.B.V.O., S.B.P., N.C.C., A.W., A.P.V., G.M.D., A-R.C. Investigation: S.B.V.O., S.B.P., N.C.C., A.W., I.B., M.M., J.X., Y.P.K., N.P.W., S.J.Y., K.M.M., J.M., S.Z., A.P.V. Writing: S.B.V.O., S.B.P., N.C.C., A.W., A-R.C. Review and Editing: S.B.V.O., S.B.P., N.C.C., T.I., A.P.V., G.M.D., A-R.C. Supervision: A-R.C.

