## Supplementary for "On the illusion of auxotrophy: *met15Δ* yeast cells can grow on inorganic sulfur thanks to the previously uncharacterized homocysteine synthase Yll058w"

**SUPPLEMENTARY INFORMATION**

**SUPPLEMENTARY FIGURES**

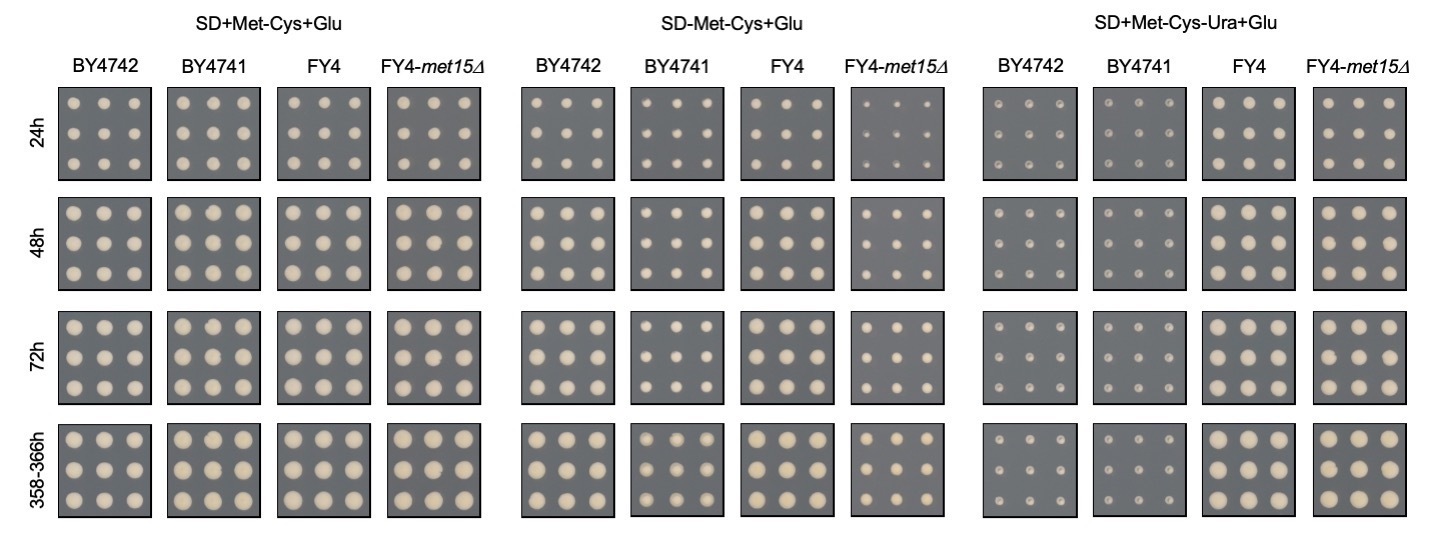

**Figure S1. Growth of *met15Δ* cells in an organosulfur-deficient medium at earlier incubation times.** In “synthetic defined” medium lacking methionine and cysteine and with glucose comprising the carbon source (SD-Met-Cys+Glu), cells harboring the *met15Δ0* deletion (BY4741) or a “scarless deletion” of *MET15* made by CRISPR (FY4-*met15Δ*) show surprisingly robust growth inconsistent with their presumed auxotrophy. Colony crops from SD+Met-Cys+Glu are shown as a positive control, and those from SD+Met-Cys-Ura+Glu to allow for comparison with an established auxotrophy. Representative colony crops are shown for growth seen at various time points prior to and at saturation.

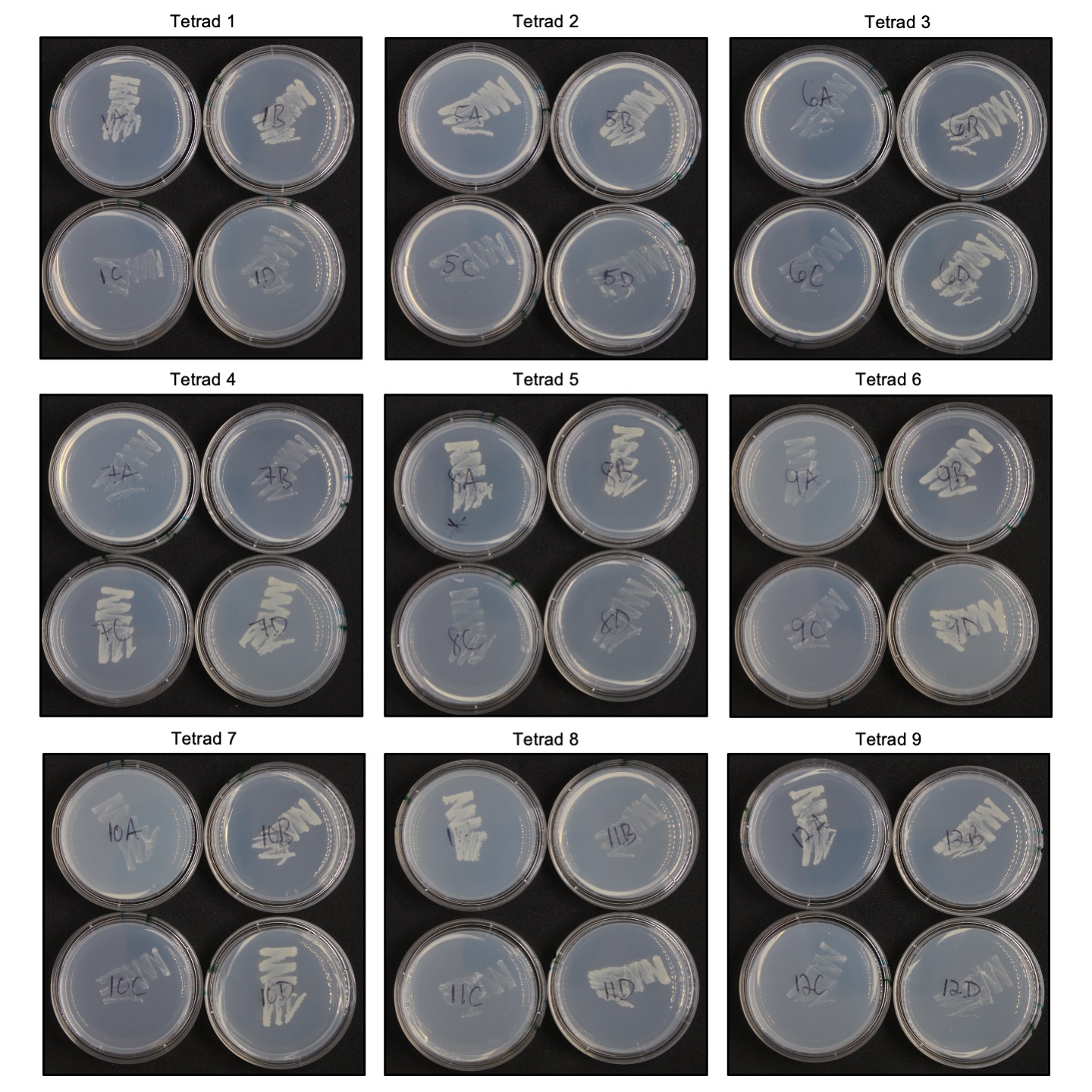

**Figure S2. *met15Δ*-associated growth phenotype segregates 2:2 in a heterozygous cross.** The BY4741 and BY4742 strains were mated, sporulated, and dissected using standard techniques. Nine, four-spore tetrads (individual panels) were grown on YPDA, and then patched onto SD-Met-Cys+Glu medium. Individual plates were used to prevent cross-feeding between *MET15+* and *met15Δ* cells. Each tetrad resulted in 2:2 segregation of the growth phenotype on organosulfur-deficient media. Notably, the growth of *met15Δ* thick patches was substantially weaker than that observed in the automated colony transfer assays used throughout the manuscript Images shown here were taken following ~72 hours of incubation at 30°C.

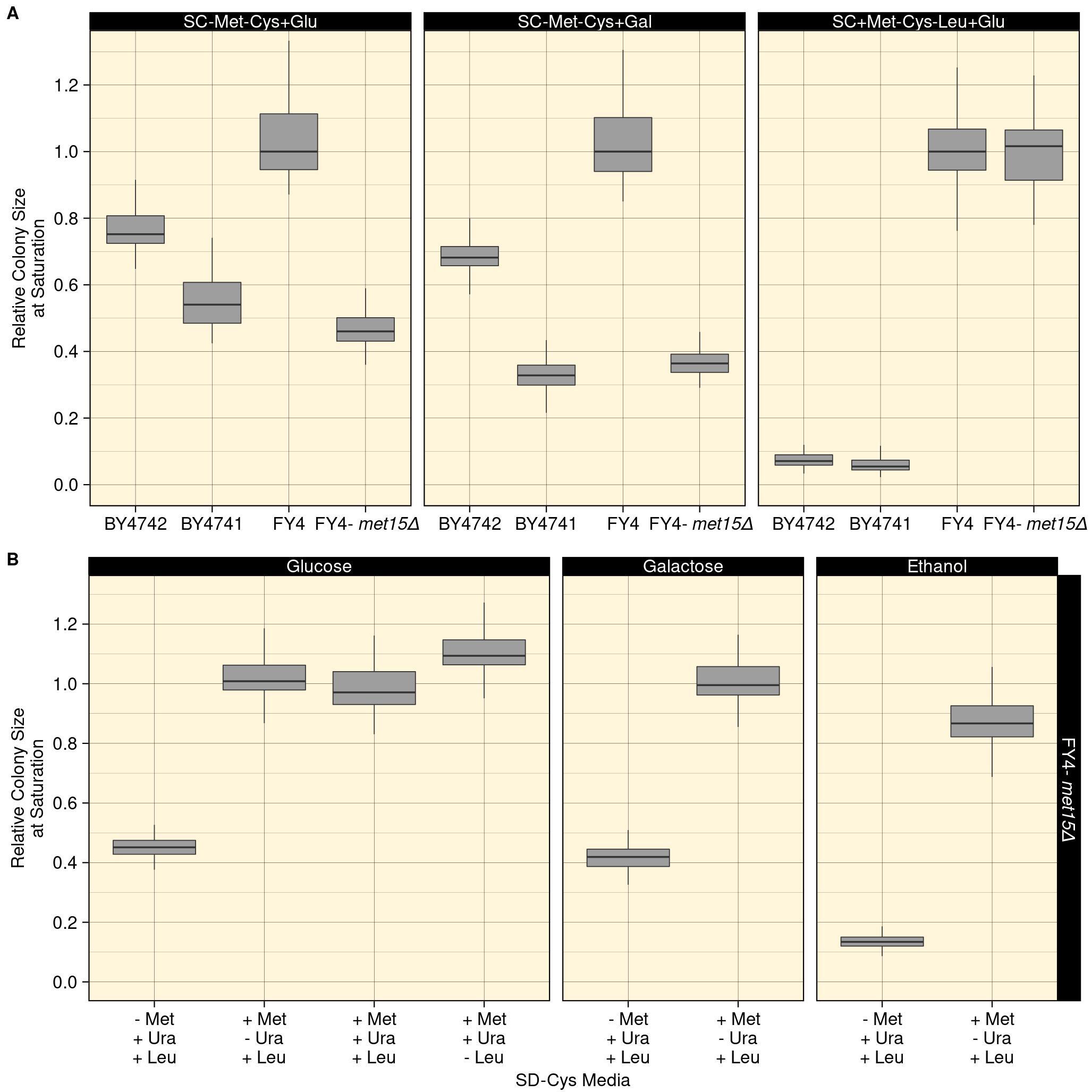

**Figure S3. Robust growth of *met15Δ* cells in media lacking organosulfurs.** **A)** The same automated colony transfer and analysis pipeline shown in **Figure 1A-C** was repeated, here on SC-Met-Cys media containing either glucose or galactose as the sole carbon source, or on SC+Met-Cys-Leu+Glu medium, where the negligible growth of the BY4741 and BY4742 strains is consistent with auxotrophy. The prototrophic strain FY4 and a mutant derivative lacking *MET15* were also assayed. Box plots represent relative colony sizes (y-axis) at saturation (y-axis, 140 hours for SC+Met-Cys-Leu+Glu and 165 hours for SC-Met-Cys+Glu/Gal). Relative colony size is normalized to the mean colony size of FY4 grown in the same condition (see **Experimental Procedures**). **B)** The “scarless” CRISPR *met15Δ* mutant shows growth in SD-Met-Cys media, inconsistent with auxotrophy, when glucose or galactose comprise the carbon source, but growth is much weaker with ethanol as the carbon source. Box plots represent relative colony sizes (y-axis) at saturation (~358 to 366 hours). Relative colony size is normalized to the mean colony size of FY4 grown in the same condition.

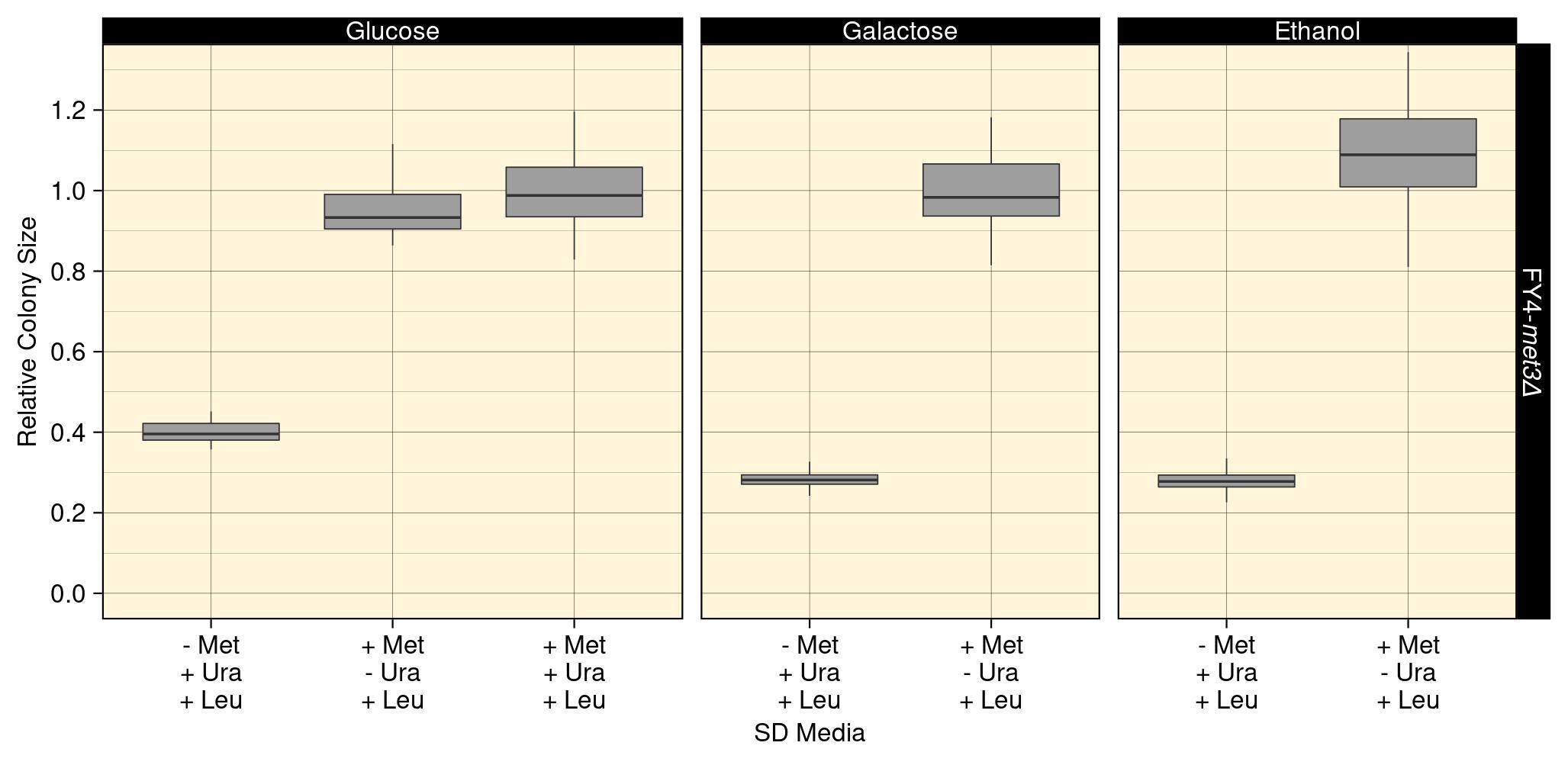

**Figure S4. The growth of FY4-*met3Δ* cells on SD-Met-Cys media is generally unaffected by the carbon source.** The FY4-*met3Δ* mutant was grown in the same manner as the strains presented in **Figures 1C** and **S3**. Box plots represent relative colony sizes (y-axis) at saturation (366 hours). Relative colony size is normalized to the mean colony size of FY4 grown in the same condition.

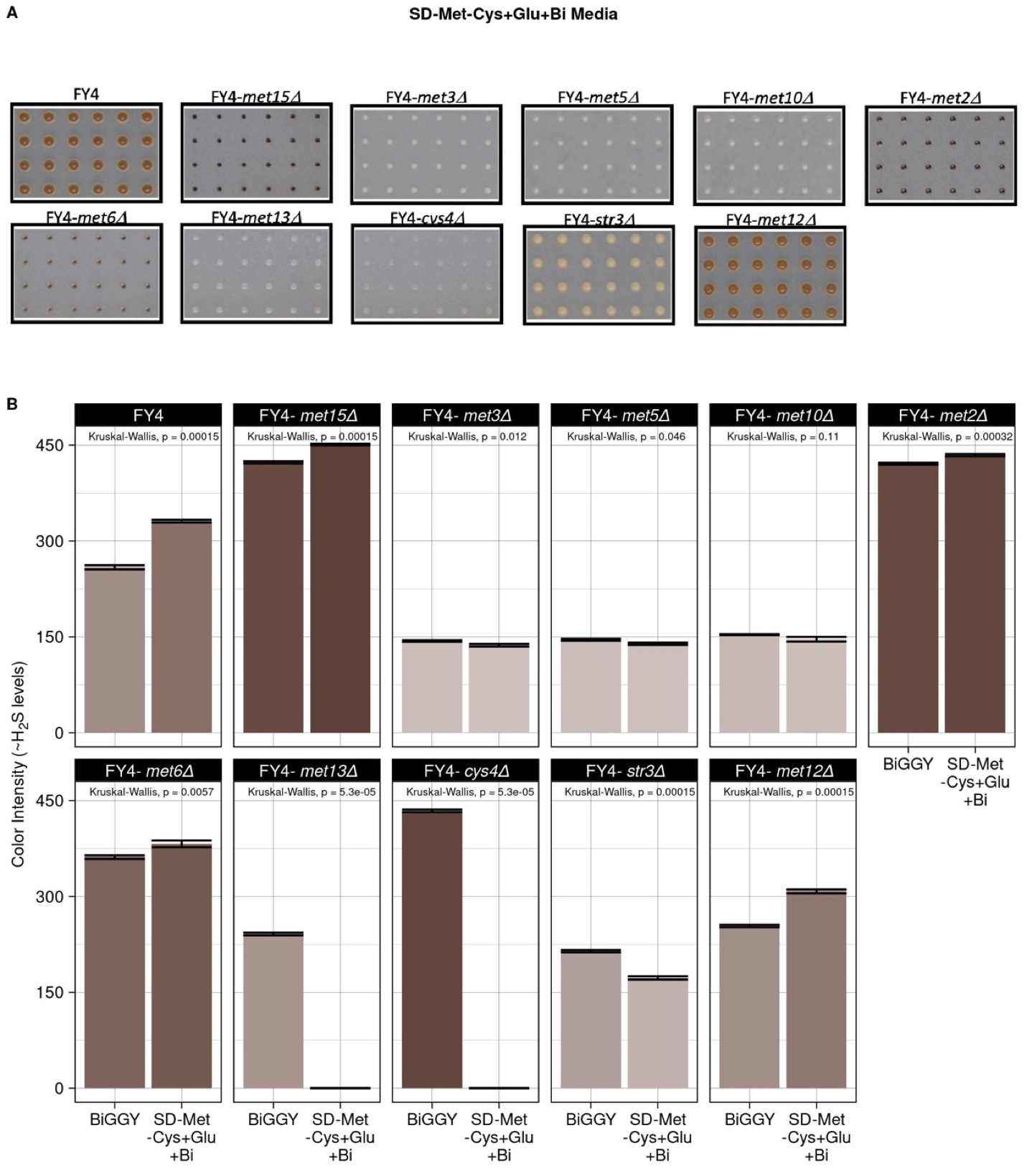

**Figure S5. Relative H_2_S levels of mutant strains.** **A)** Cropped images of colony color of mutant strains pinned to SD-Met-Cys+Glu medium containing bismuth (SD-Met-Cys+Glu+Bi) at saturation (173.5 hours). **B)** Quantitative assessment using [RGB] data (see **Supplementary Materials and Methods**), of H_2_S in the strains (individual panels) grown in rich media containing bismuth (BiGGY; see **Figure 2C**) and SD-Met-Cys+Glu+Bi (**Panel A**). Higher values correspond to darker color. Error bars: standard error. p-values on the top of the bars are from the Kruskal-Wallis comparison of the color intensity in the two media. Values in SD-Met-Cys+Glu+Bi are set as 0 for FY4-*met13Δ* and FY4-*cys4Δ* because these strains did not grow at all on this media.

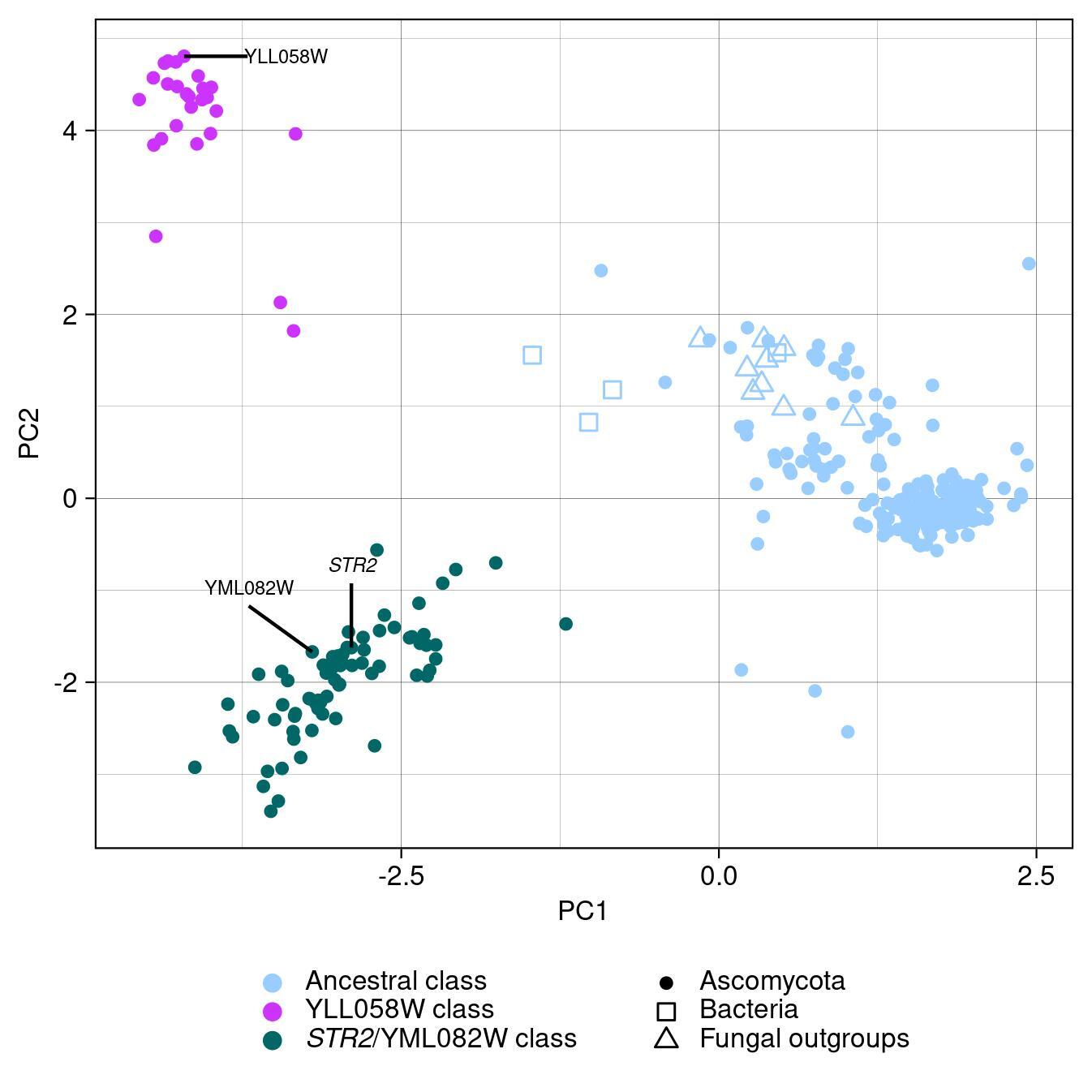

**Figure S6. YLL058W homologs cluster into three distinct classes.** The sequence distances between YLL058W homologs found in budding yeasts and outgroup taxa are visualized using multidimensional scaling (MDS). Points are colored according to k-means clustering based on these sequence distances using three clusters. The three genes contained in the *S. cerevisiae* genome are labeled.

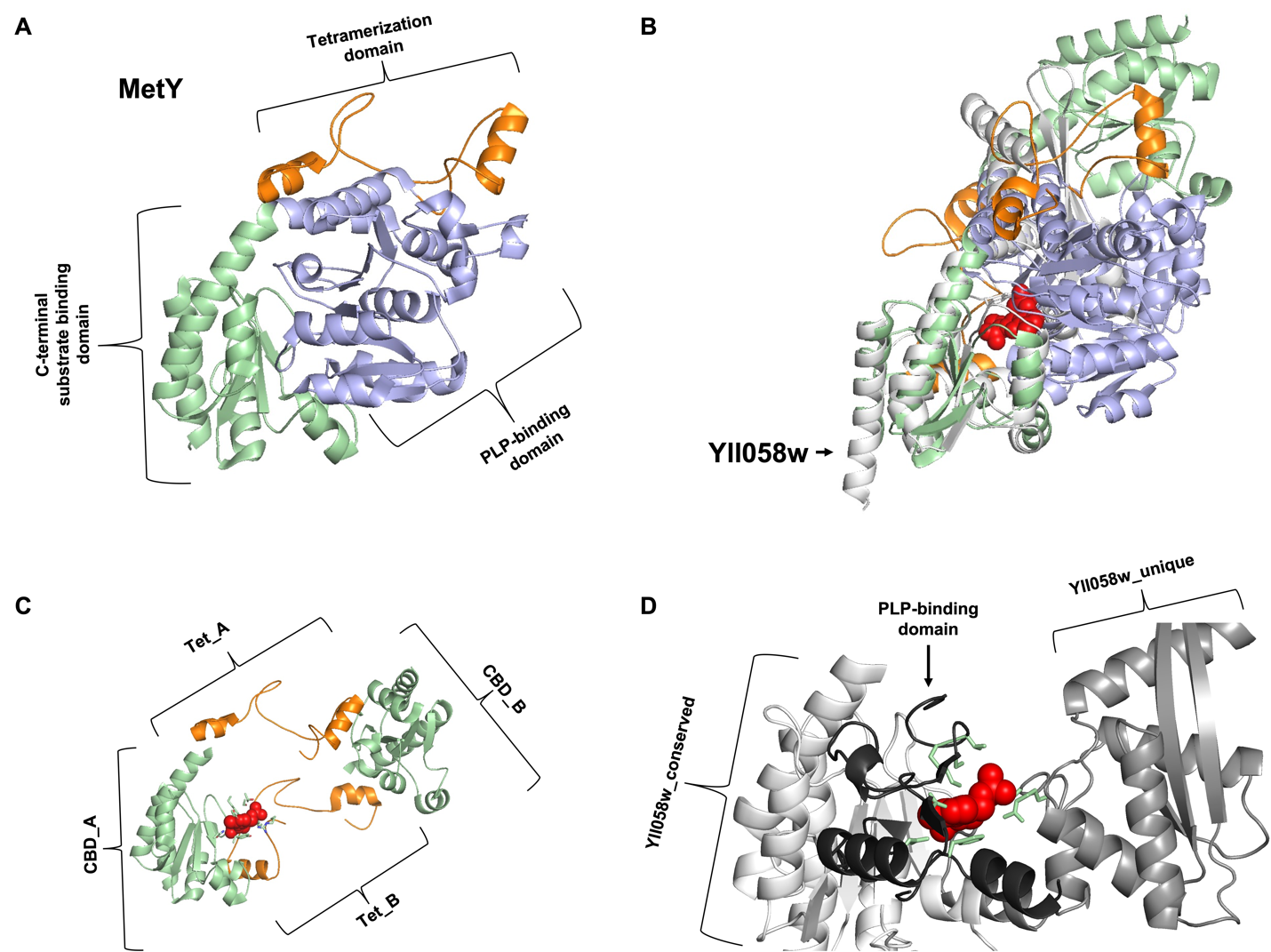

**Figure S7. Yll058w functions as a monomer, in contrast to the tetrameric conformation seen in canonical homocysteine synthases. A)** A cartoon view of the MetY monomer, from the MetY crystal structure, pdbID:7KB1 (1). The N-terminal tetramerization domain is shown in orange, the PLP-binding domain is shown in light purple, and the C-terminal substrate binding domain is shown in pale green. **B)** The predicted structure of the monomeric Yll058w (white) is highly similar structurally to MetY within the PLP-binding and C-terminal domains and aligns well with a MetY dimer (reaction intermediate colored red for clarity). Orange, light purple and pale green indicate the same domains as in Panel A. **C)** Cartoon view of the MetY N-terminal tetramerization domains (Tet_A and Tet_B; orange) and the C-terminal substrate binding domains (CBD_A, CBD_B; pale green) in contact with the reaction intermediate (red). The PLP-binding domains are hidden. Met15 (not shown) is predicted to adopt a similar conformation. **D)** The unique portion of the Yll058w monomer supplies the additional catalytic residues (pale green) provided by the second subunit of the MetY dimer. The portion of the predicted structure of Yll058w in contact with the reaction intermediate (red) is shown, with areas showing sequence and predicted structural similarity shown in white, and the region lacking homology to MetY and Met15 (Yll058w_unique) shown in gray. This unique region supports the possibility of Yll058w catalysis as a monomer, as opposed the confirmed (MetY) and predicted (Met15) multimeric structures of other homocysteine synthases. Portions of the PLP-binding domain that contribute putative catalytic residues are shown in black.

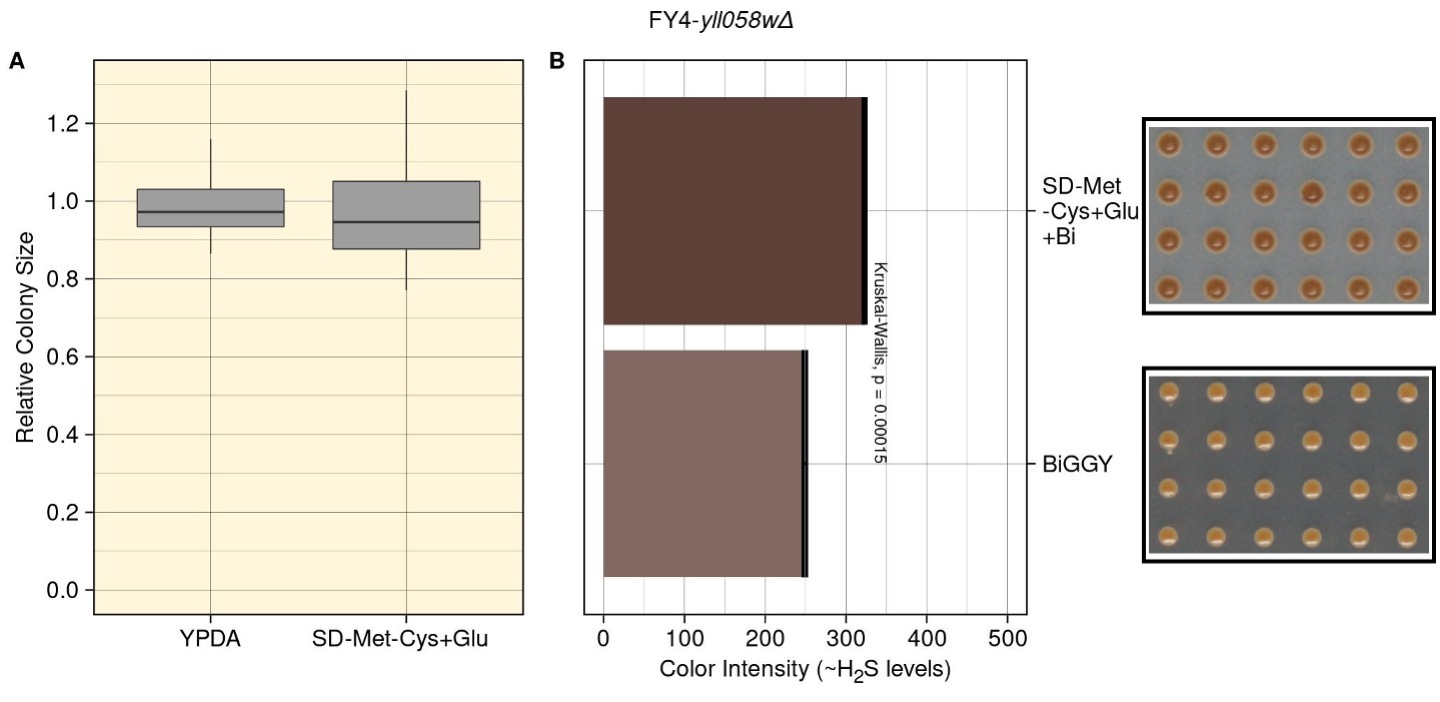

**Figure S8. Deletion of YLL058W has negligible effect on fitness and H_2_S accumulation in the FY4 background. A**) Deletion of YLL058W in the prototrophic FY4 background has a negligible effect on growth in SD-Met-Cys+Glu medium. The strains were examined in the same manner as the mutants shown in **Figure 2B**, with colony size normalized to that of FY4. **B)** The relative H_2_S level of FY4*-yll058wΔ*, as determined by color when growing on BiGGY and SD-Met-Cys+Glu+Bi, is comparable to that of wild-type FY4 cells. The darkness of the colonies was determined using the same method applied to the mutants in **Figure S5B.**

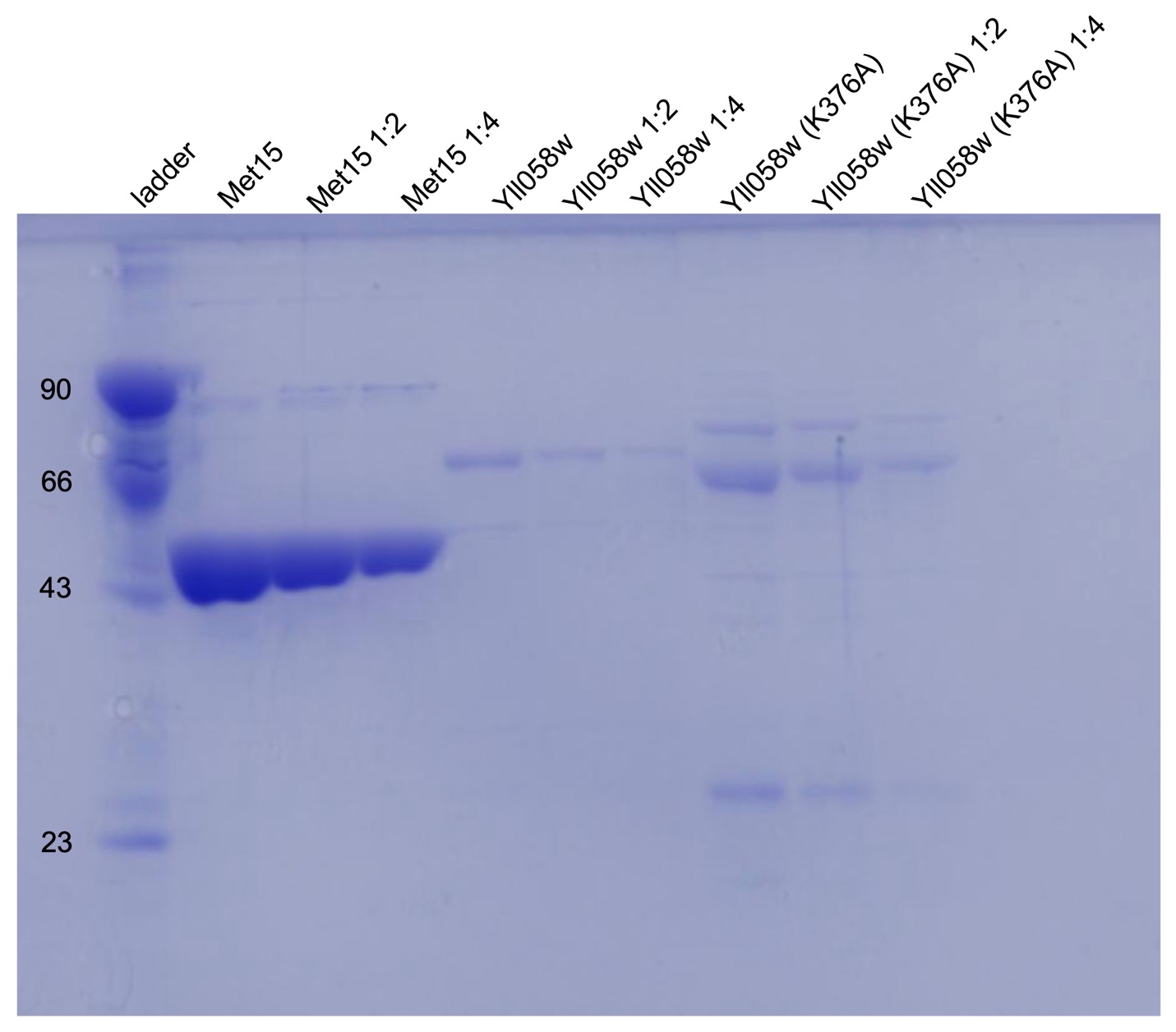

**Figure S9. Coomassie-stained SDS-PAGE gel showing the final fractions that contained the recombinantly-expressed, purified protein used in the *in vitro* homocysteine biosynthesis assay.** A lane containing molecular weight standards (kDa = kilodaltons) is shown on the left. The molecular weights of Met15, Yll058w and Yll058w (K376A) are 48.66, 64.23 and 64.17 kDa respectively. The K376A mutation in Yll058w likely affects protein folding and thereby affects the speed at which the protein runs through the gel, resulting in a slightly higher band than expected. The other two non-specific bands in the Yll058w (K376A) lanes are impurities.

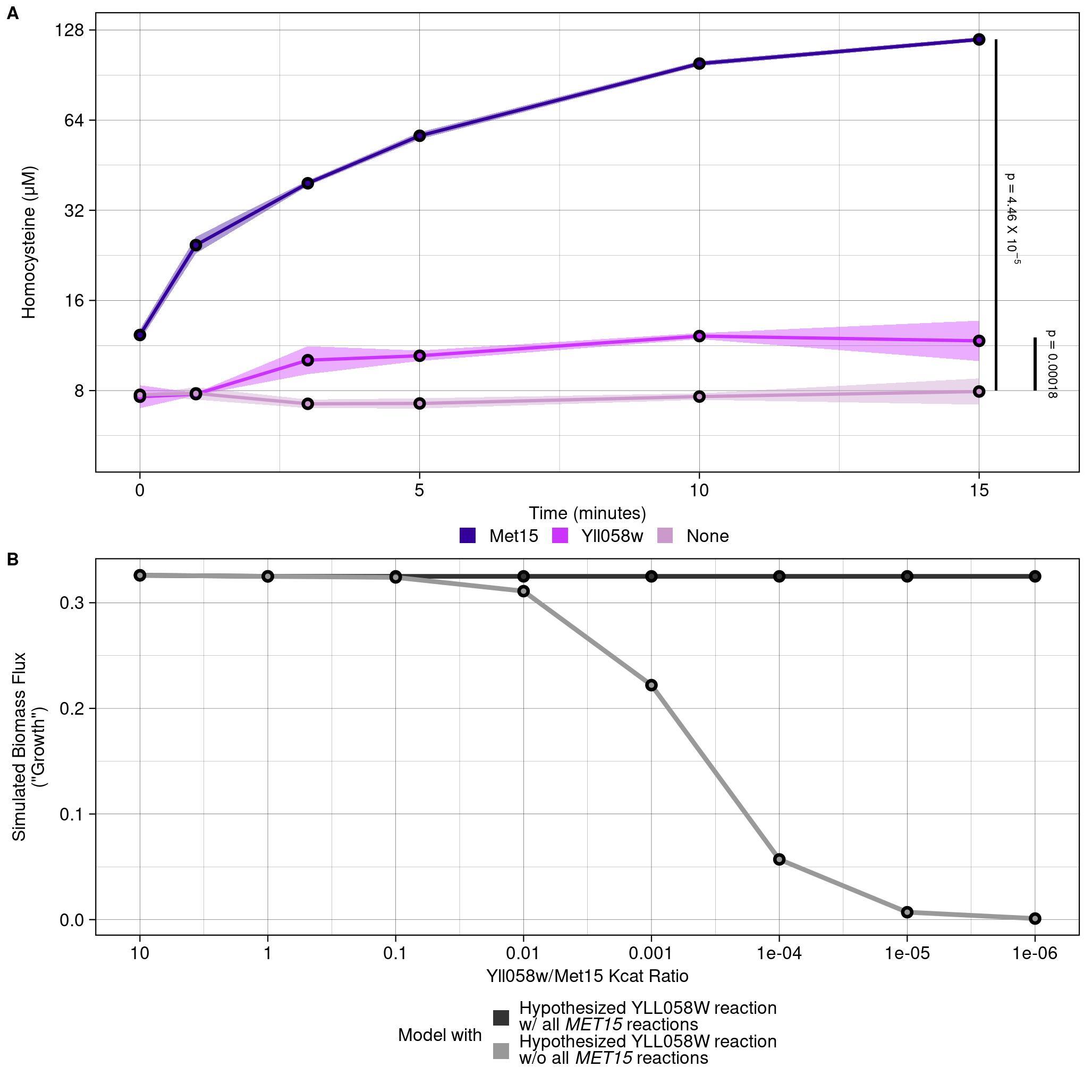

**Figure S10.** **Yll058w catalyzes homocysteine biosynthesis but less efficient than Met15**. **A)** An in vitro homocysteine biosynthesis assay was carried out using recombinantly expressed, purified *S. cerevisiae* Met15 (violet) or Yll058w (magenta), or a no-enzyme control (pink), as the catalyst. Homocysteine levels (y-axis; log2 scale) are shown over time (x-axis). Solid lines: mean. Shaded area: one standard deviation. p: two-tailed, paired Student’s T-test paired combining all replicates and time points, comparing homocysteine levels for Met15 and Yll058w to the negative control. **B)** Growth in an organosulfur-free medium is predicted *in silico* even with a homocysteine synthase with a dramatically lower catalytic efficiency than Met15. Two modifications were made to the default model – the addition of the *in vitro* reaction carried out in **Panel A** utilizing Yll058w as the catalytic enzyme (black), and the subsequent removal of the *MET15* gene from the model (gray). The y-axis indicates the flux through the “biomass equation”, a proxy for growth/fitness, and the x-axis indicates different catalytic efficiency (Kcat) values assigned to Yll058w, relative to the default Kcat assigned to Met15, with the efficiency of Yll058w in the hypothetical reaction decreasing from left to right.

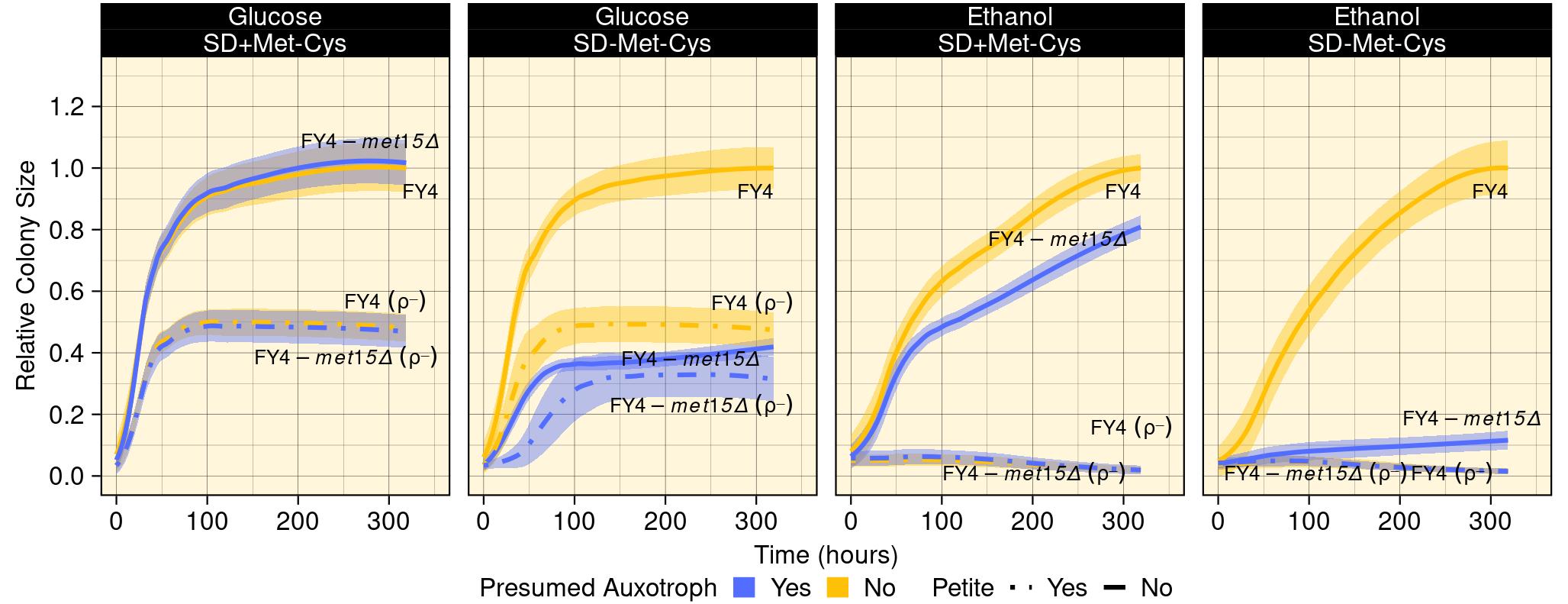

**Figure S11. A growth defect of *met15Δ* cells grown with ethanol is seen in the presence and absence of organosulfurs in the media.** Growth curves of FY4 and the FY4-*met15Δ* mutant (solid line), along with their respective petite (ρ^-^) derivatives (dashed line), grown on SD media containing or lacking methionine and utilizing either glucose or ethanol as the sole carbon source are shown. For each condition (subpanel) the cells were grown as described in **Figure 1A and** **Experimental Procedures**. All colony sizes were normalized to the median colony size of FY4 at the final time point in the same condition. Line: mean relative colony size. Shaded regions: one standard deviation. **Table S5** contains the Kruskal-Wallis Test results comparing the growth of a particular strain as estimated by area under the curve in the presence or absence of methionine in media containing either glucose or ethanol as the carbon source.

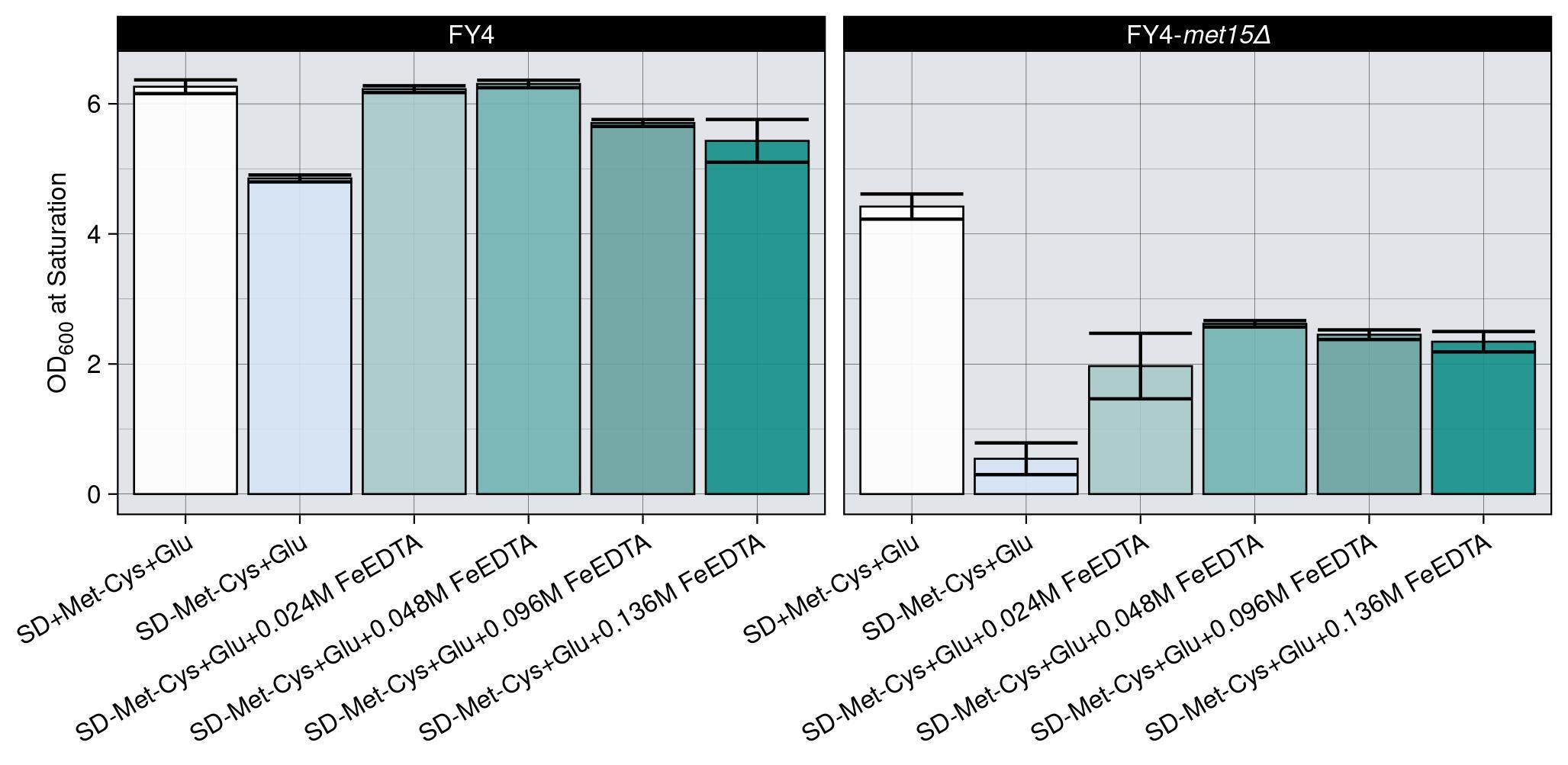

**Figure S12. Titration of the H_2_S chelator Fe-EDTA.** To determine the optimal dosage of Fe-EDTA for the experiments depicted in **Figures 4B-D**, FY4 and FY4-*met15Δ* cultures were grown in SD+Met-Cys+Glu and SD-Met-Cys+Glu supplemented with Fe-EDTA, with concentrations raging from 0 to 0.136 M. The concentration of 0.048 M yielded the greatest restoration of growth in FY4-*met15Δ* cultures. Bar plot: mean OD_600_ at saturation (~51 hours). Error bars: standard error.

**SUPPLEMENTARY TABLES**

**Table S1.** Settings for the RoToR HDA robotic plate handler used for all the automated pinning experiments described in the manuscript.

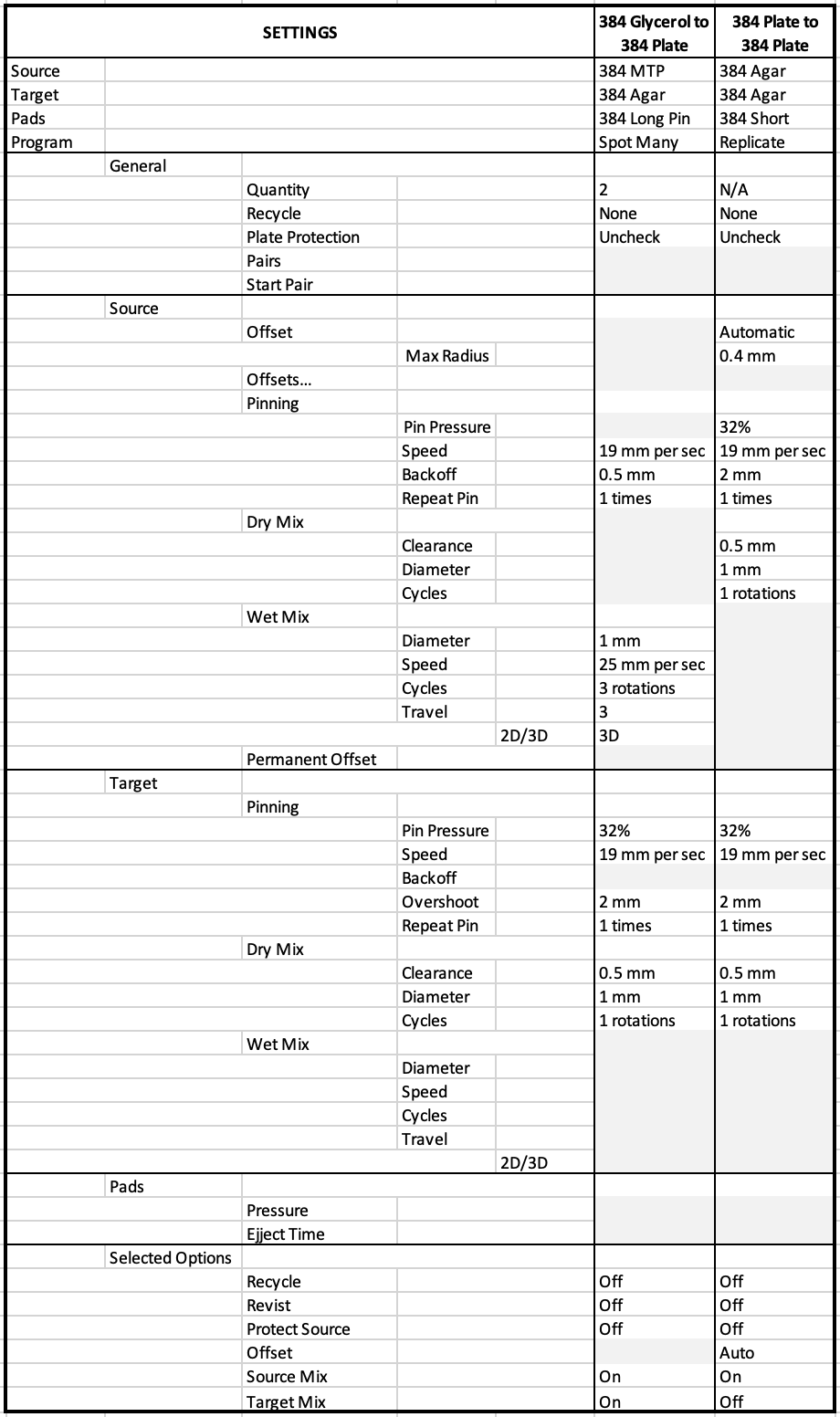

**Table S2.** Mass spectrometry analysis confirms that the growth medium is not contaminated with organosulfurs. ND= Not Detected. See **Supplementary Materials and Methods**.

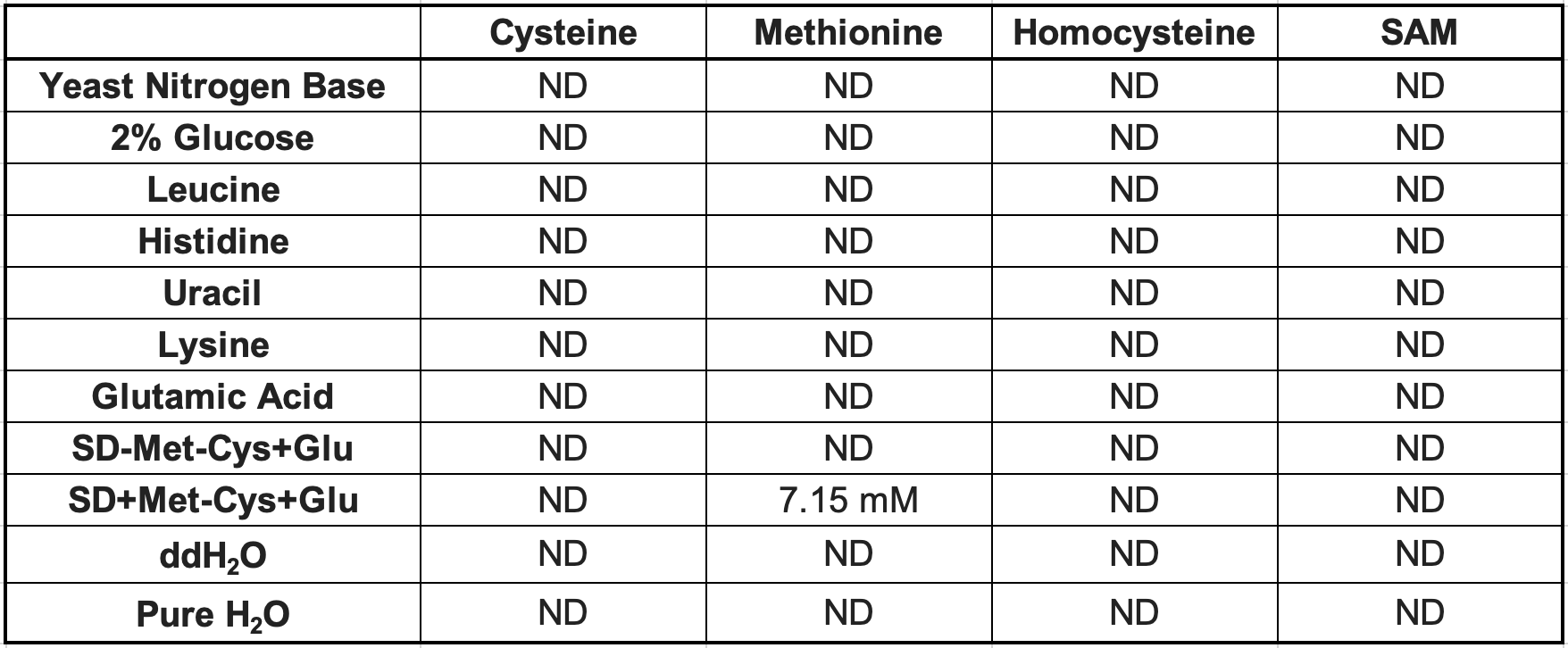

**Table S3.** YLL058W and its orthologs are found in a cluster of genes near the telomere. *S. cerevisiae* gene names are used for all species shown. The ten nearest genes (5 upstream (5’) genes, numbered -1 to -5, and 5 downstream (3’) genes, numbered 1 to 5) to YLL058W orthologs are shown. Asterisks indicate a known role in sulfur metabolism.

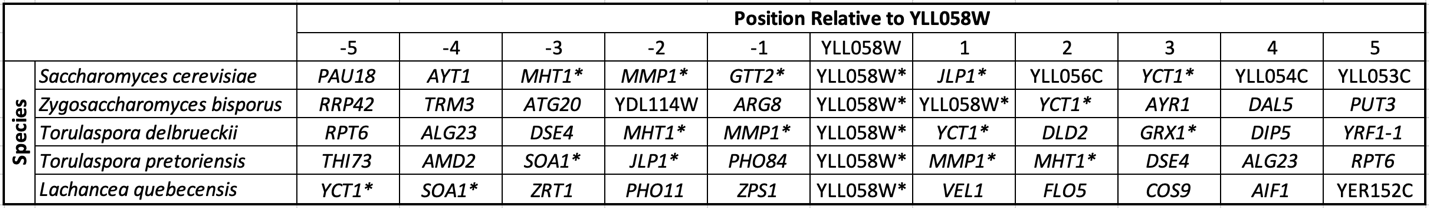

**Table S4.** OD_600_ values at saturation (~51 hours) for cultures of the indicated strains grown in SD-Met-Cys+Glu medium with or without the H_2_S chelator Ferric(Fe)-EDTA (column 4), and OD_600_ values for the same cultures diluted and repassaged in SD-Met-Cys+Glu medium lacking chelator (column 5). Each strain has 3 replicates (1-3) from two independent experiments (A&B). Although FY4-*met15Δ* cultures generally failed to show any substantial growth in liquid medium without an H_2_S chelator present (**Figure 4B**), we occasionally observed “escape” in four FY4-*met15Δ* cultures (indicated in green text) grown in SD-Met-Cys+Glu without an H_2_S chelator present, independent of whether the cells had previously been in the presence of chelator.

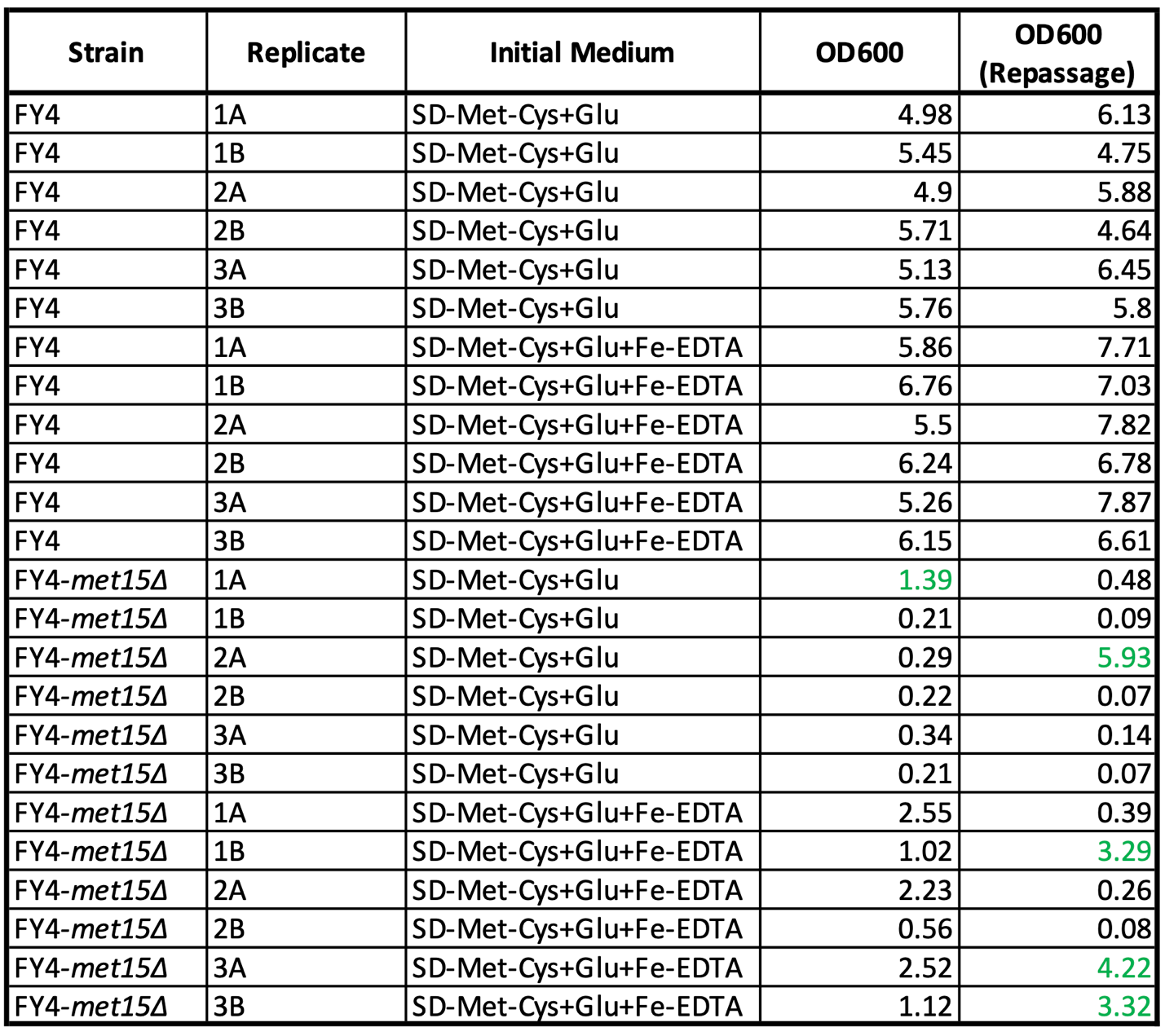

**Table S5.** Kruskal-Wallis Test results comparing the growth of a given strain in condition_1 (+Met) vs. condition_2 (-Met), with EtOH or Glu. The statistic and p-value columns are the Kruskal-Wallis test statistic and p-value output, and the effect size column is the difference between the median fitness value in media lacking methionine and that in media containing methionine divided by the median fitness value in media containing methionine.

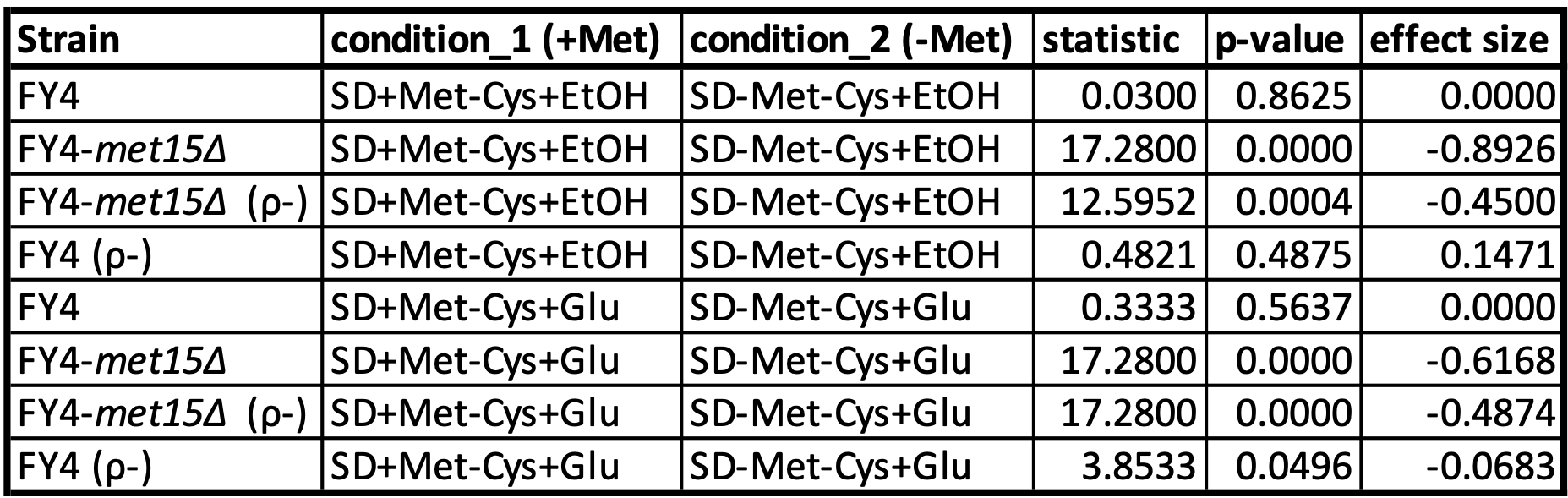

**Table S6.** *Saccharomyces cerevisiae* strains used in this study.

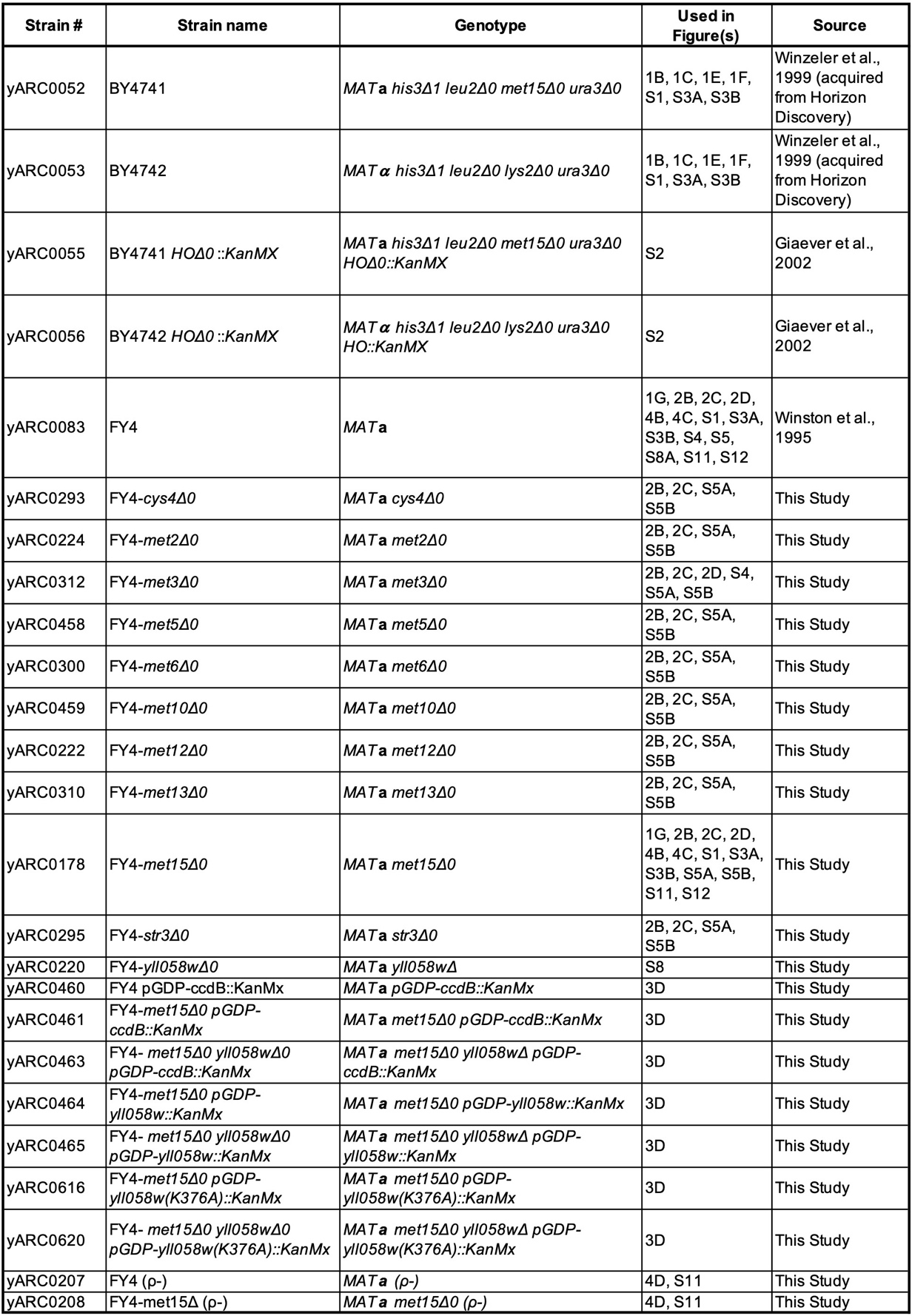

**Table S7.** Plasmids used in this study.

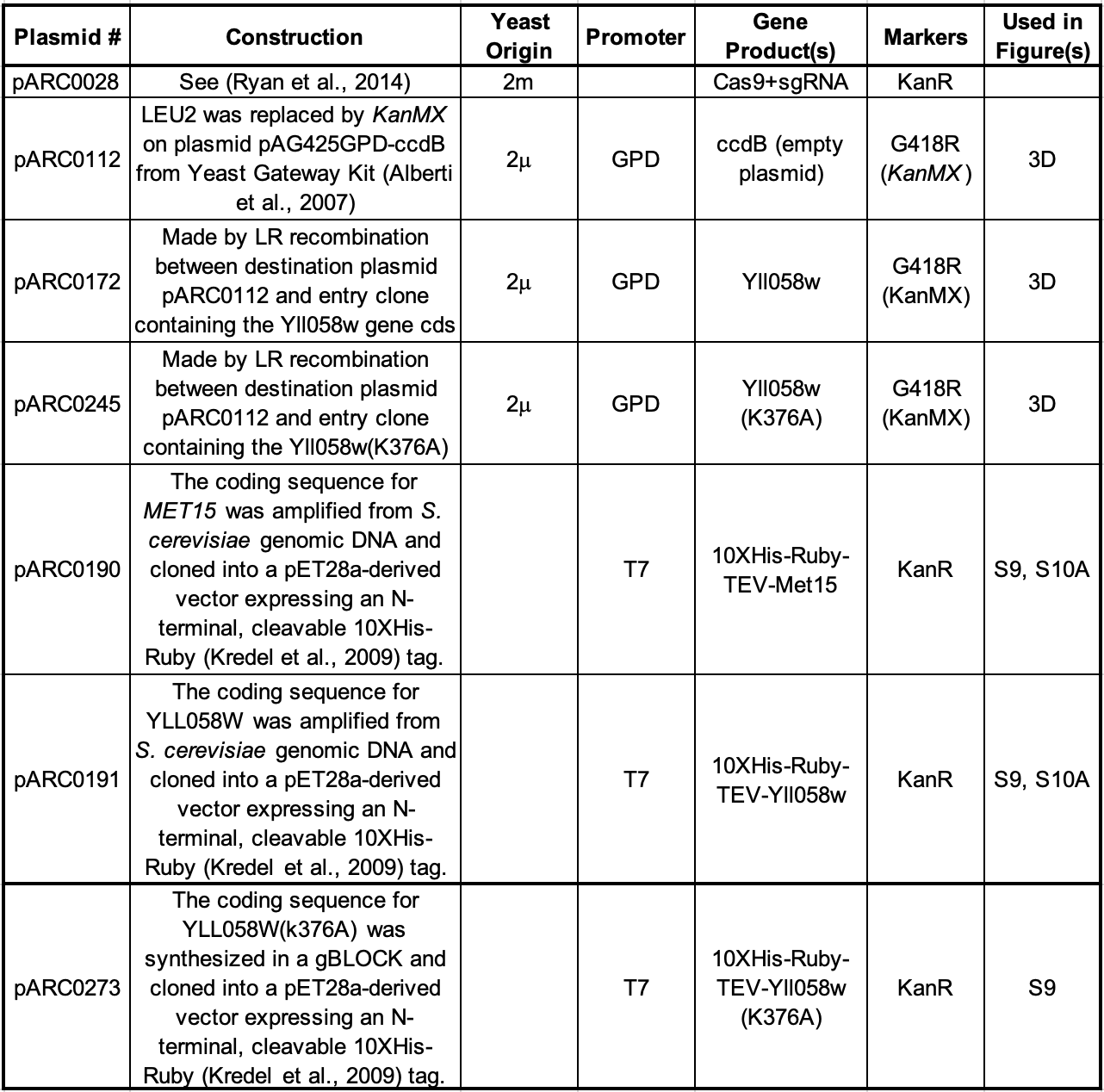

**Table S8.** sgRNA, repair fragment template (HR_template) and amplification primer (HR_Fw and HR_rv) sequences used for CRISPR/Cas9-mediated ORF deletions.

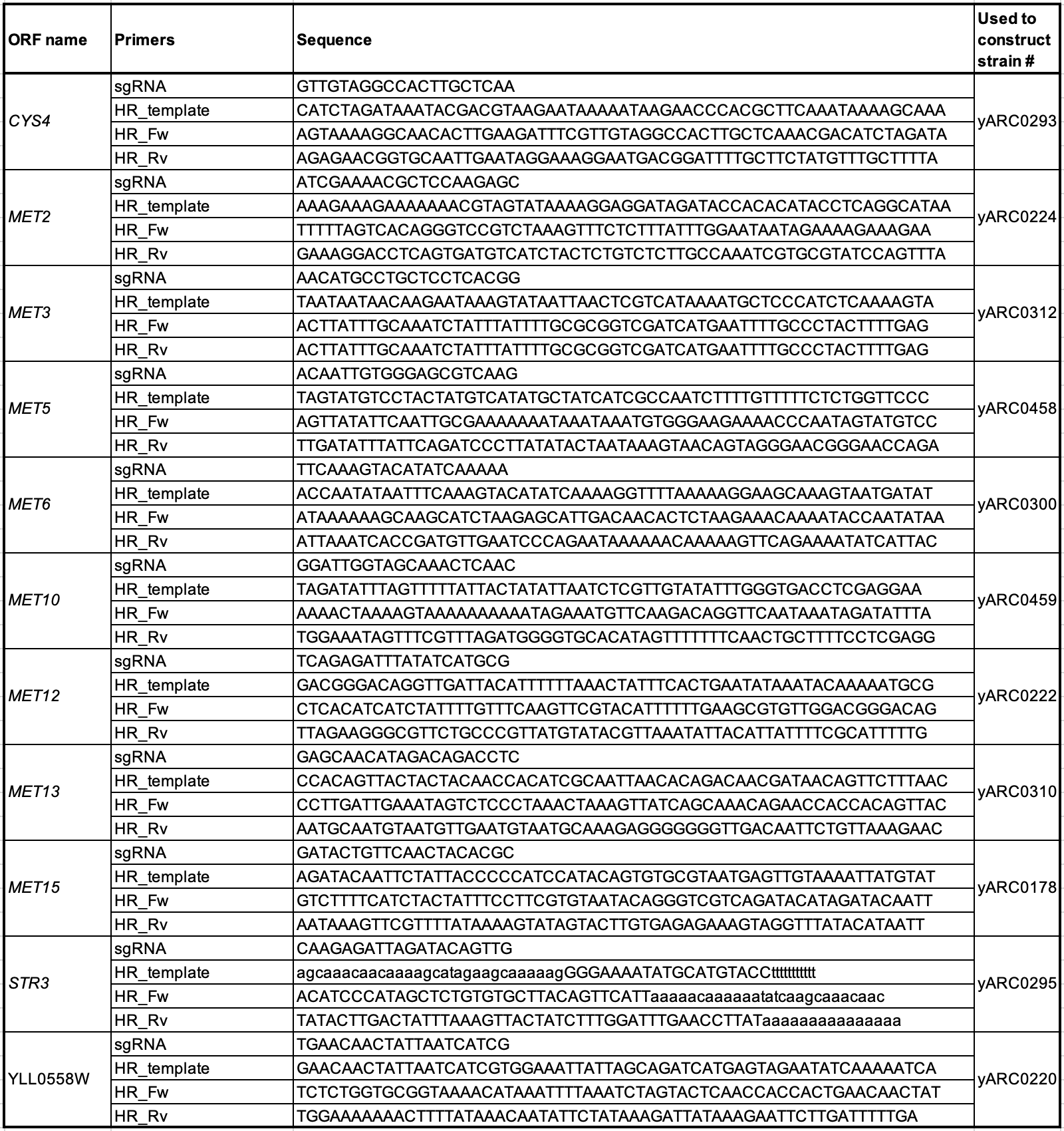

**Table S9.** Yeast growth conditions in this study. When required, the analogous liquid medium was made with the same composition, excluding the 2% agar.

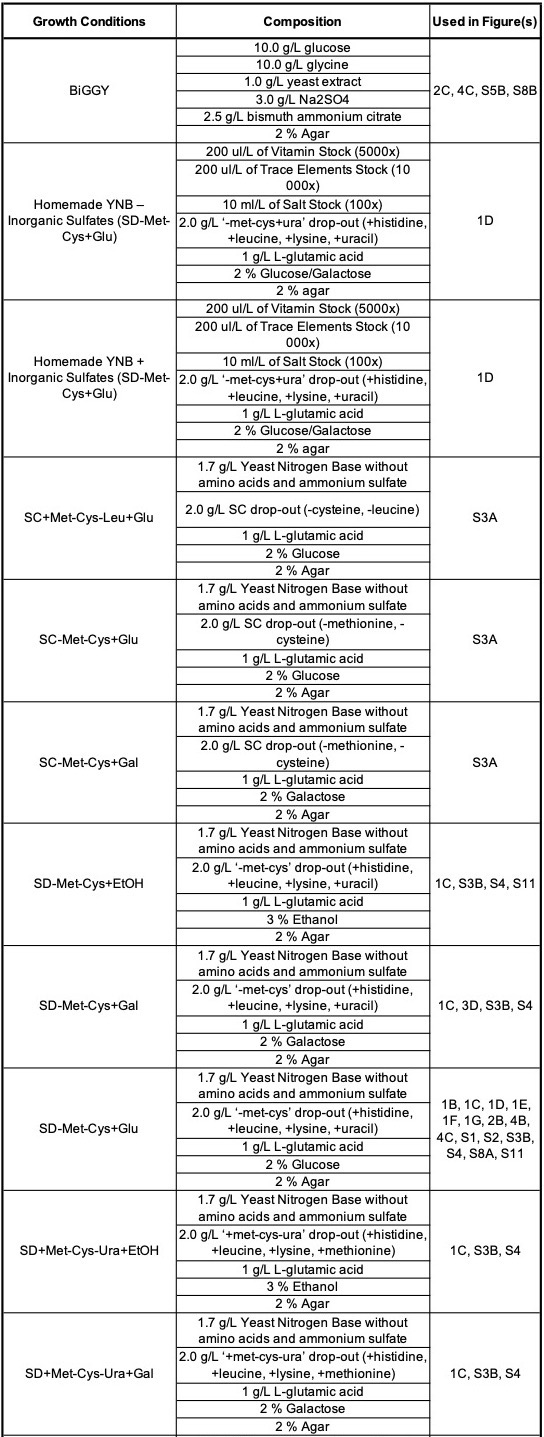

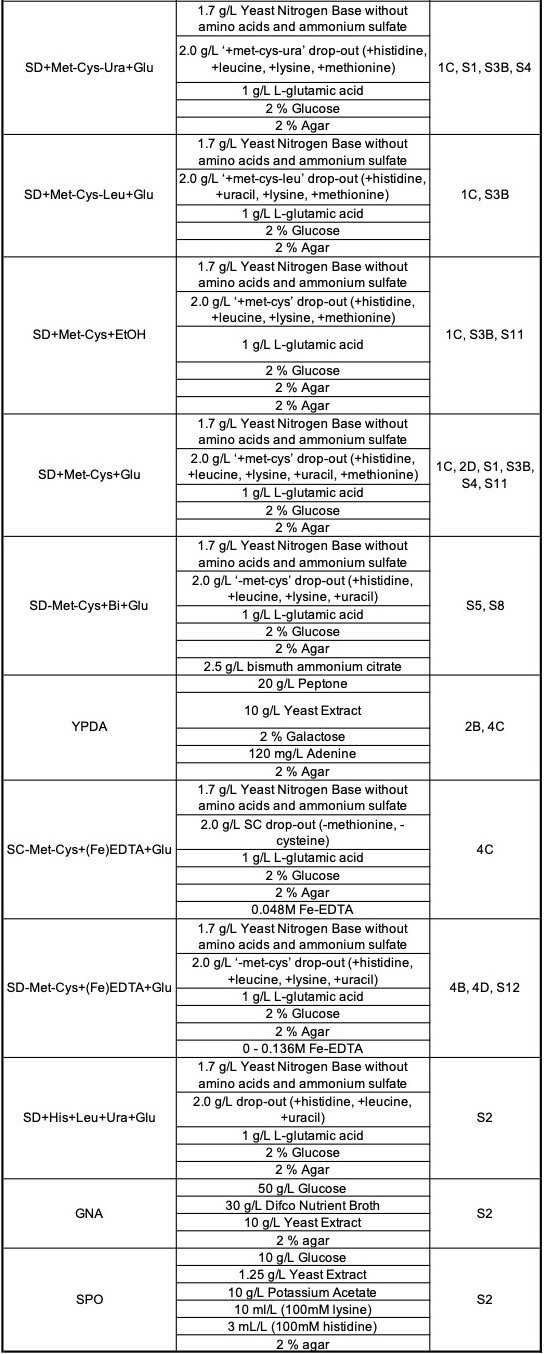

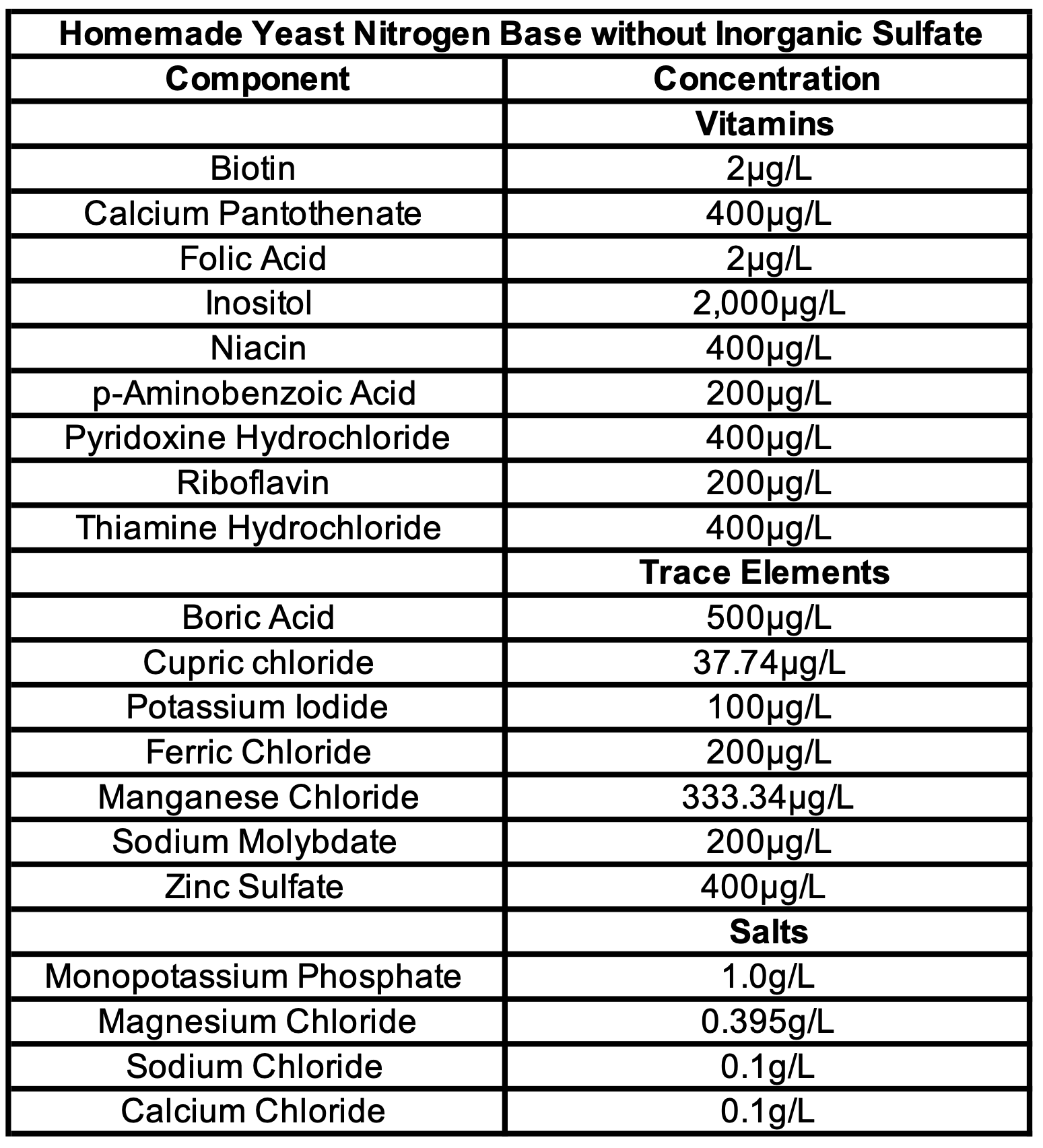

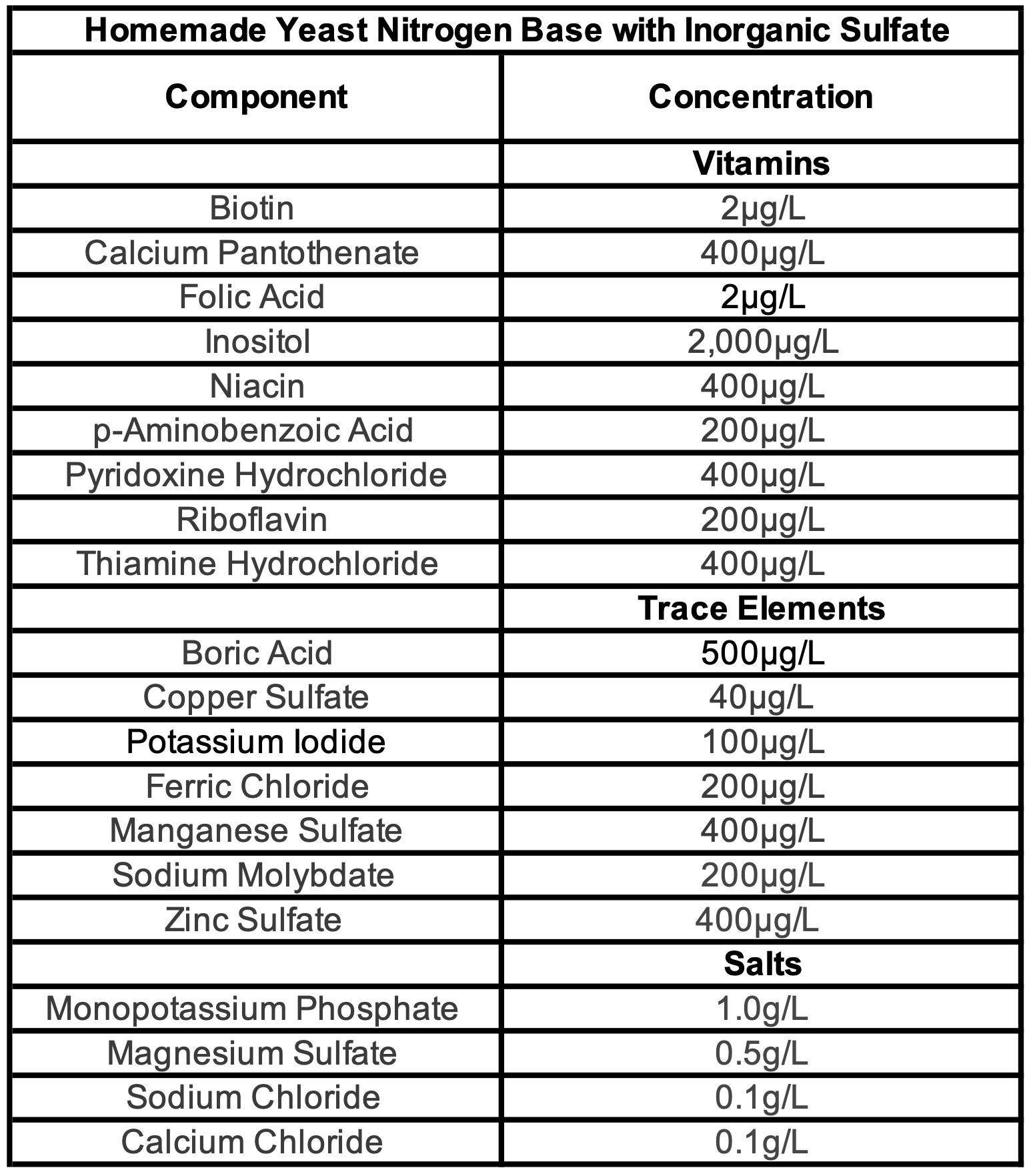

**Table S10.** K_m_ and V_max_ values for Met15, Yll058w and Yll058w (K376A)

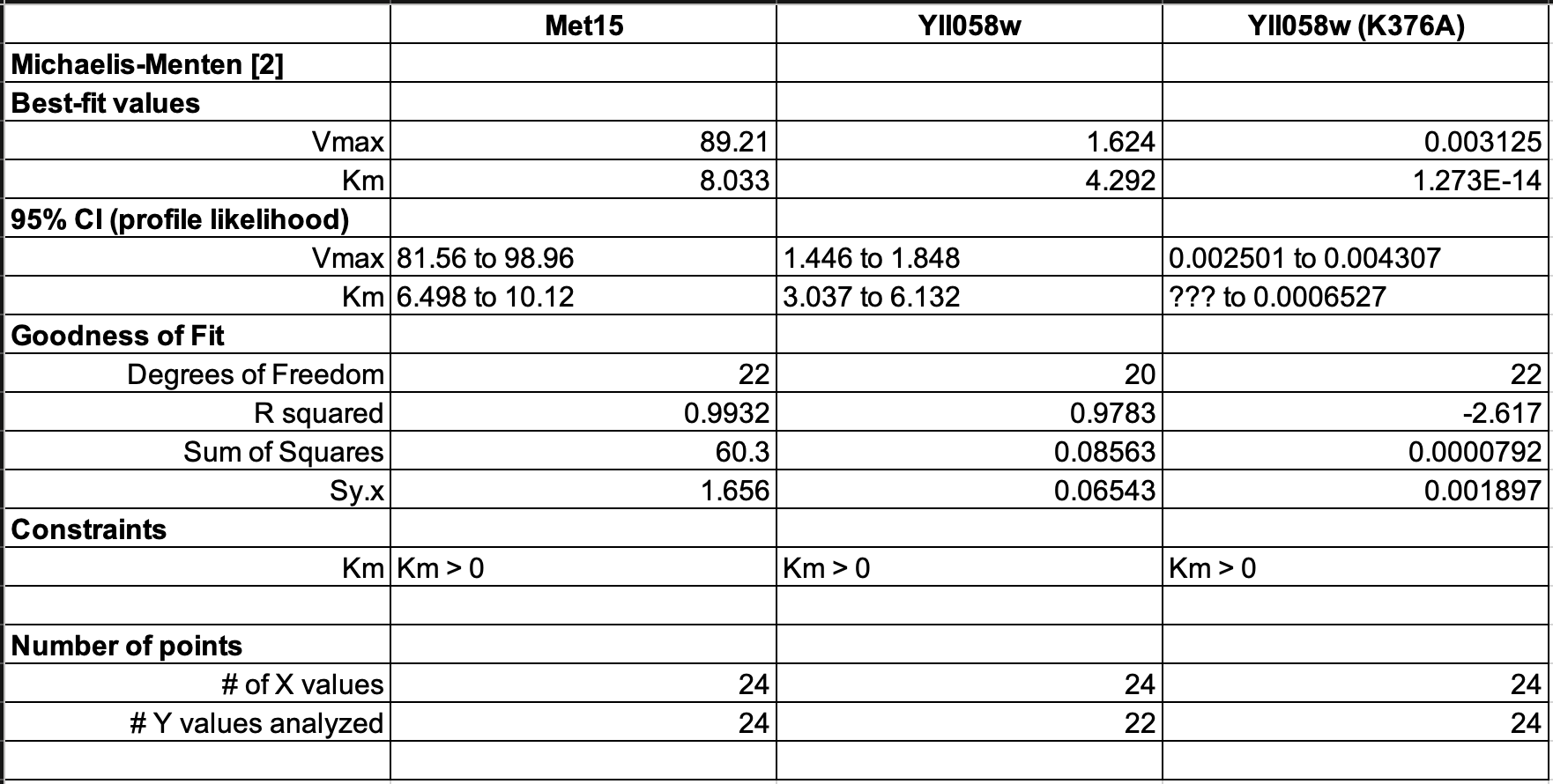

**SUPPLEMENTARY MATERIALS AND METHODS**

**Gene deletion with CRISPR/Cas9**

Co-transformations were made with 5μg of repair fragment and 1μg of plasmid expressing cas9 (Addgene 60847, **Table S7**) (6) with the respective sgRNA (**Table S8**) (9). Identification of PAM sites and sgRNA selection were chosen using Benchling (<https://www.benchling.com/>). Transformed cells were plated on YPDA+G418 and colonies were PCR genotyped to select those with the correct gene deletion. The positive clones were then grown on YPDA to induce the loss of the cas9- and sgRNA-containing plasmids. Strains carrying plasmids for overexpression experiments were made with the LiAc/PEG/ssDNA protocol (10) using 500ng of each plasmid and selected on YPDA+G418.

**Construction of petite strains**

Petite derivatives of FY4 and FY4-*met15Δ* were made using a slightly modified version of the standard LiAc/PEG/ssDNA yeast transformation protocol (10), with shorter incubation times and without a DNA template, followed by replica platting on YPDA and YP-Glycerol plates to identify the petites.

**Plasmid construction**

The overexpression plasmids used in **Figure 3D** were made by replacing the Leu2 gene by KanMx cassette into the destination plasmid pAG425GPD-ccdB (7). This plasmid was used as an empty vector control (pARC0112, **Table S7**) while an LR recombination (Gateway LR Clonase II Enzyme Mix, ThermoFisher) of this same plasmid with an entry clone containing either the ORF YLL058W or YLL058W (K376A) were used to produce the plasmids overexpressing YLL058W and YLL058W (K376A) (pARC0172 and pARC0245, respectively, **Table S7**). Plasmids pARC0190 and pARC0191 (**Table S7**) were made by amplification of *MET15* and YLL058W coding sequences, respectively, from FY4 genomic DNA, and cloned into a pET28a-derived vector expressing an N-terminal, cleavable 10XHis-Ruby tag (8). The plasmid pARC0273 was done following the same strategy, but using a synthetic gBlock of the YLL058W sequence, containing the K376A modification.

**Strain Crossing**

The cross depicted in **Figure S2** was carried out by standard techniques. Briefly, the BY4741 and BY4742 deletion collection parent strains (**Table S6**) were mated on YPDA solid medium and incubated at room temperature (RT) for ~6 hours, and then struck to select for colonies formed from single diploids on solid SD+His+Leu+Ura medium (**Table S9**). After incubation at 30°C for ~2 days on the diploid selection medium, colonies were patched to “GNA” plates (**Table S9**) and incubated at 30°C for ~2 days. These plates were then replica plated to “sporulation (SPO)” medium (**Table S9)** containing histidine, lysine, methionine, uracil, and leucine, and incubated at RT for ~11 days. Four-spore tetrads were selected and dissected using standard techniques and initially grown on YPDA. Individual spores were then patched to petri dishes (**Figure S2**).

**Colony size estimation**
Serial imaging of plates was done either using spImager Automated Imaging System (S & P Robotics Inc., Ontario, Canada) or manually using a custom-made lightbox with an overhead camera mount and a commercially available SLR camera (18Mpixel Rebel T6, Canon USA Inc., Melville, NY, USA). Plates were imaged at regular intervals beginning right after transfer until the colonies reached saturation. Images were analyzed in bulk using a custom script made using functions from the MATLAB Colony Analyzer Toolkit (11) to provide colony size estimations (<https://github.com/sauriiiin/lid_personal/blob/master/justanalyze.m>). Output files containing colony size information along with the images are available at <https://bit.ly/3pOe6aT>.

**Relative H_2_S measurement**

Bismuth containing plates (BiGGY and SD-Met-Cys+Glu+Bi; **Table S9**) were used to assess the level of excess sulfides in various strains. Cells were transferred from YPDA plates onto bismuth containing plates, as described on **Figure 1A**. Color intensity was measured using the pixel intensity values from the darkest regions of the colonies. To do this we made use of the Digital Color Meter application in MacOS (Version 13.0 Beta (22A5321d)) with the ‘display native values’, and lowest aperture size option. We then selected five colonies per replicate (two) per strain and measured the red and green pixel intensity within the darkest region of the colony. The sum of the red and green pixel intensity value was subtracted from 510 (255*2, the maximum possible pixel intensity in the red and green pixels) and used as the ‘color intensity’. We had 10 color intensity measurements per strain per media. The darker the color of the colony the higher the color intensity.

**Mass spectrometry analysis of media by semi-targeted high-resolution LC-HRMS**

**Sample preparation**

Metabolic quenching and polar metabolite pool extraction was performed by adding 400µL ice cold methanol to 100µL of sample. Deuterated (D_3_)-creatinine and (D_3_)-alanine, (D_4_)-taurine and (D_3_)-lactate (Sigma-Aldrich) was added to the sample lysates as an internal standard for a final concentration of 10µM. Samples were vortexed, and then homogenized using a 25°C water bath sonicator for 5 minutes. The supernatant was then cleared of protein by centrifugation at 16,000xg. Two µL of cleared supernatant was subjected to online LC-MS analysis.

**LC-HRMS Method**

Analyses were performed by untargeted LC-HRMS. Briefly, Samples were injected via a Thermo Vanquish UHPLC and separated over a reversed phase Thermo HyperCarb porous graphite column (2.1×100mm, 3μm particle size) maintained at 55°C. For the 20 minute LC gradient, the mobile phase consisted of the following: solvent A (water / 0.1% formic acid) and solvent B (ACN / 0.1% formic acid). The gradient was the following: 0-1min 1% B, increase to 15%B over 5 minutes, continue increasing to 98%B over 5 minutes, hold at 98%B for five minutes, reequillibrate at 1%B for five minutes. The Thermo IDX tribrid mass spectrometer was operated in positive ion mode, scanning in ddMS^2^ mode (2 μscans) from 70 to 800 m/z at 120,000 resolution with an AGC target of 2e5 for full scan, 2e4 for ms^2^ scans using HCD fragmentation at stepped 15,35,50 collision energies. Source ionization setting was 3.0kV spray voltage for positive mode. Source gas parameters were 35 sheath gas, 12 auxiliary gas at 320°C, and 8 sweep gas. Calibration was performed prior to analysis using the Pierce^TM^ FlexMix Ion Calibration Solutions (Thermo Fisher Scientific). Integrated peak areas were then extracted manually using Quan Browser (Thermo Fisher Xcalibur ver. 2.7) (Metabolomics & Lipidomics Core, University of Pittsburgh Health Sciences Core Research Facilities).

**Purification of recombinantly expressed proteins**

Met15, Yll058w and Yll058w (K376A) were expressed as fusion proteins with an N-terminal His_10_-mRuby2 tag which can be removed by cleavage with TEV protease. Fusion proteins were expressed in the BL21 (DE3) RIPL CodonPlus *E. coli* strain (Stratagene). Cells were grown at room temperature in 2L of standard LB media containing kanamycin and chloramphenicol to an OD600 of ~0.6, induced with 0.5mM IPTG, and grown for eight additional hours at room temperature post-induction. Purification of both proteins was performed essentially as described previously (12). Briefly, all proteins were purified by two rounds (pre- and post-cleavage with TEV protease) of nickel affinity chromatography followed by ion exchange chromatography. Met15 was further purified by size exclusion chromatography. The final purified fractions that were used in the *in vitro* homocysteine assay are shown in **Figure S9**.

**Measurement of homocysteine levels from *in vitro* reactions**

Ten µL of each enzyme reaction were extracted and derivatized with N-ethylmaleamide (NEM**; Alfa Aesar, Cat# 40526-06**) in 990 µL of ice-cold extraction solvent (80% MeOH: 20% H_2_O containing 25 mM NEM and 10 mM ammonium formate, pH=7.0) containing 0.5 uM of [D_4_]-Homocysteine **(Cambridge Isotope labs, Cat# DLM-8259-PK)** followed by incubation at 4°C for 30 min. After centrifugation (17,000 g, 20 min, 4°C), the supernatants were analyzed by LC-MS following previously established conditions (13). For the chromatographic metabolite separation, a Vanquish UPLC system was coupled to a Q Exactive HF (QE-HF) mass spectrometer equipped with HESI (Thermo Fisher Scientific, Waltham, MA). The column was a SeQuant ZIC-pHILIC LC column, 5 mm, 150 x 4.6mm (MilliporeSigma, Burlington, MA) with a SeQuant ZIC-pHILIC guard column, 20 x 4.6 mm (MilliporeSigma, Burlington, MA). Mobile phase A was 10 mM (NH_4_)_2_CO_3_ and 0.05% NH_4_OH in H_2_O while mobile phase B was 100% ACN. The column chamber temperature was set to 30°C. The mobile phase condition was set according to the following gradient: 0-13min: 80% to 20% of mobile phase B, 13-15min: 20% of mobile phase B. The ESI ionization mode was positive. The MS scan range (m/z) was set to 60-900. The mass resolution was 120,000 and the AGC target was 3 x 10^6^. Capillary voltage and capillary temperature were set to 3.5 KV and 320°C, respectively. Five μL of sample was loaded. The homocysteine and [D_4_]-Homocysteine peaks were manually identified and integrated with EL-Maven (Version 0.11.0) by matching with an in-house library. Homocysteine levels were calculated using the standard curve and corrected using the [D_4_]-Homocysteine internal standard.
